# Adaptive behavior is guided by integrated representations of controlled and non-controlled information

**DOI:** 10.1101/2025.08.07.669231

**Authors:** Bingfang Huang, Harrison Ritz, Jiefeng Jiang

## Abstract

Understanding how task knowledge is encoded neurally is crucial for uncovering the mechanisms underlying adaptive behavior. Here, we test the theory that all task information is integrated into a conjunctive task representation by investigating whether this representation simultaneously includes two types of associations that can guide behavior: stimulus-response (non-controlled) associations and stimulus-control (controlled) associations that inform how task focus should be adjusted to achieve goal-directed behavior. We extended the classic item-specific proportion congruency paradigm to dissociate the electroencephalographic (EEG) representations of controlled and non-controlled associations. Behavioral data replicated previous findings of association-driven adaptive behaviors. Decoding analyses of EEG data further showed that associations of controlled and non-controlled information were represented concurrently and differentially. Brain-behavioral analyses also showed that the strength of both associations was associated with faster responses. These findings provide initial evidence supporting the idea that controlled and non-controlled associations are governed by an integrated task representation to guide adaptive behaviors simultaneously.

## Introduction

Cognitive control coordinates our thoughts and actions with internal goals (Egner, 2017; Miller & Cohen, 2001) via the abilities of sustained task focus when faced with distractions (stability) and flexible adjustments between task sets based on varying situations (flexibility). Both stability and flexibility can be implemented in a proactive (Botvinick et al., 2001; Jiang et al., 2015) or reactive (Blais et al., 2007; Verguts & Notebaert, 2008) manner (Abrahamse et al., 2016; Braem & Egner, 2018; Braver, 2012; Brown et al., 2007; Egner, 2014, 2023; Jiang et al., 2018). For task focus indicating cognitive stability, conflict paradigms have been employed to explore how the brain focuses on task-relevant information while avoiding the distraction from potent task-irrelevant information (Eriksen & Eriksen, 1974; Simon & Rudell, 1967; Stroop, 1935). For example, in the Stroop task (Stroop, 1935), participants are shown a color word and asked to respond to the ink color of the word while ignoring the meaning of the word. The ink color can be consistent (congruent trials) or inconsistent (incongruent trials) with the meaning of the word, resulting in a conflict effect measured by worse performance on incongruent trials than on congruent trials. Some models of cognitive control postulate that the conflict effect reflects the involvement of cognitive control, which suppresses the habitual but inappropriate response to the word and/or boosts the novel and goal-directed response to the ink color to resolve the conflict on incongruent trials (Botvinick et al., 2001; Cohen et al., 1990).

A key question in the research on cognitive control is how the brain monitors control demands (i.e., how much control is needed). Recent research has demonstrated the importance of associative learning in guiding control demand (Bustamante et al., 2021; Jiang et al., 2020a; Jiang et al., 2020b; Wen et al., 2023). In particular, theories focusing on stimulus-control (SC) associations assume that the brain binds together a stimulus feature with the concurrent control demands (Bejjani et al., 2020; Blais et al., 2007; Bugg & Crump, 2012; Bugg & Hutchison, 2013; Bugg et al., 2011; Chiu et al., 2017; Colvett et al., 2025; Ileri-Tayar et al., 2024; Ileri-Tayar et al., 2022; Ileri-Tayar et al., 2025; Jiang et al., 2020a; Spinelli et al., 2022; Suh & Bugg, 2021). For example, in the Stroop task, if the color red is used on mostly congruent (MC) trials and the color blue is presented on mostly incongruent (MI) trials, then red would be associated with lower control demands relative to blue. This process has been proposed to account for the item-specific proportion congruency (ISPC) effect (Braem et al., 2019; Jacoby et al., 2003), whereby the conflict effect is reduced for stimulus features that predict higher control demands.

The ISPC effect is a natural prediction of stimulus-control learning, with the potential to provide a theoretical foundation for how cognitive control is scaffolded on structured representations of our environment. However, there have been theoretical challenges that the ISPC may arise from lower-level stimulus-response (SR) learning theory instead. This alternative account posits that associations may also form between stimulus and specific response (e.g., the most-likely response) in addition to more abstract SC associations (Atalay & Misirlisoy, 2012; Hazeltine & Mordkoff, 2014; Schmidt & Besner, 2008; Schmidt et al., 2016). Returning to the Stroop task, if the word blue is presented in red color (i.e., an incongruent trial), then the word blue will be associated with the ‘red’ response (in Stroop, often an assigned keypress or verbal response). Note that a stimulus may be associated with multiple responses. For example, if the word blue is presented in blue color (i.e., a congruent trial), then the word blue will also be associated with the ‘blue’ response. In this case, SR association is linked to the most-likely response. Following the example above, if the word blue is presented mostly on incongruent trials, then the word blue is paired more often with the red response than the blue response, and the SR is between the word blue and the red response. Subsequent responses to this stimulus will be biased towards the associated response, whether or not they are correct.

Since SC and SR accounts often predict similar behavioral patterns, previous studies have typically focused on one at a time through experimental controls, and current empirical evidence supports both SC (Bugg & Hutchison, 2013; Gonthier et al., 2016; Spinelli & Lupker, 2020a; Spinelli et al., 2019) and SR (Schmidt & Besner, 2008) associations. Additionally, studies of ISPC have differentiated the behavioral effects and neural substrates of SC and SR associations when either SC or SR associations are available (Bugg et al., 2011; Chiu et al., 2017). For example, Chiu and colleagues (Chiu et al., 2017) showed that the dynamics of SC and SR learning were captured by trial-wise prediction errors with a reinforcement learning model and that SC learning was encoded in the caudate nucleus while SR association was represented in the parietal cortex. Conceptually, SC associations affect behavior by modulating other cognitive processes via cognitive control, whereas SR associations drive behavior in a non-controlled manner by retrieving the associated response. These accounts provide competing explanations for what kind of knowledge is encoded in neural task representations. Task representations are often thought to be conjunctive, providing one-point access to all task-related information (Badre, 2024; Badre et al., 2021; Hommel, 2004, 2019; Kikumoto et al., 2024; Kikumoto & Mayr, 2020; Kikumoto et al., 2022a; Kikumoto et al., 2022b; Sakai, 2008; Schumacher & Hazeltine, 2016a; Verguts & Notebaert, 2008). However, it remains unexplored whether and how controlled and non-controlled information can be integrated into a task representation to guide adaptive behavior. If controlled and non-controlled information are encoded in the same representation, then: (1) SC and SR representations should occur simultaneously; (2) the strength of SC and SR representations, being in the same task presentation, should covary (i.e., positively correlated) over time as the strength of the task representation fluctuates and (3) both SC and SR associations should exhibit behavioral relevance.

To test these predictions, we extended the classic ISPC paradigm to dissociate each of SC and SR associations from their associated stimuli (e.g., separating SC association from the stimulus and the linked control demand). Using this new ISPC paradigm in an EEG experiment, we tested whether SC and SR form integrated task representations using decoding, representational subspace analysis, and representational similarity analysis (RSA). We found three main results on task representations that integrate controlled (SC) and non-controlled (SR) associations. First, time-resolved RSA on the decoding results showed simultaneous representations of SC and SR associations. Second, the strength of SC and SR associations was positively correlated, consistent with the hypothesis that both SC and SR associations are integrated in a task representation. Third, both the strength of SC and SR associations were associated with faster responses at the trial level, indicating the behavioral relevance of the neural representations. These results provide the initial evidence of an integrated task representation that contains both SC and SR associations to jointly guide goal-directed behaviors.

## Results

### Task overview

EEG data were acquired while participants (N = 40) performed a 4-key Stroop task. On each trial, participants responded to the color of a word while ignoring the identity of the word (e.g., RED presented in blue color; Fig. 1a). Participants were encouraged to respond as quickly and accurately as possible. We applied three key manipulations to this experiment (Fig. 1b). First, each stimulus could be either congruent or incongruent, defined by whether the color was consistent with the word. Second, to test whether controlled and non-controlled information are encoded in a task representation, we manipulated the ISPC (Bugg et al., 2011; Bustos et al., 2024; Chiu et al., 2017; Jacoby et al., 2003) in Phase 1 to allow the associations between colors and their paired cognitive control demands (stimulus-control; SC) and the associations between words and their most-likely response (i.e., not necessarily the actual response on the current trial; stimulus-response; SR) to be induced at the same time. To keep the overall proportion of congruency at 50%, one set of colors was presented on 75% of congruent trials (mostly congruent trials, or MC), whereas the other set of colors was presented on 25% of congruent trials (mostly incongruent trials, or MI). The MC manipulation pairs color with low control state based on SC assumption (e.g., red-congruent) and word with the consistent response (e.g., red-red) based on the SR account. Similarly, the MI manipulation links color to high control state (e.g., blue-incongruent) and word to the inconsistent response (e.g., yellow-green, because each set has two colors, the inconsistent response is unique). This standard ISPC manipulation can test whether neural representations of controlled and non-controlled information are wrapped on the same trial by combing with the following EEG analysis (See Methods). However, it potentially mixes color identity with SC, word identity with SR, and the ISPC between SC and SR. To de-confound these factors when estimating SC and SR association representations on each trial, we modified this paradigm by flipping the ISPC contingencies across different phases of the task. For example, the color ‘red’ might be associated with 25% congruency in Phase 1, 75% congruency in Phase 2, and then return to 25% congruency in Phase 3 (see Methods). The color groups were counterbalanced across participants by red and blue as the color set of MC in one group while as the color set of MI in another group in the phase 1. Overall, these manipulations resulted in a 2 (Congruency: congruent vs. incongruent) × 2 (ISPC: MC vs. MI) × 3 (Phase 1 - 3) within-subject factorial design.

**Figure 1.**
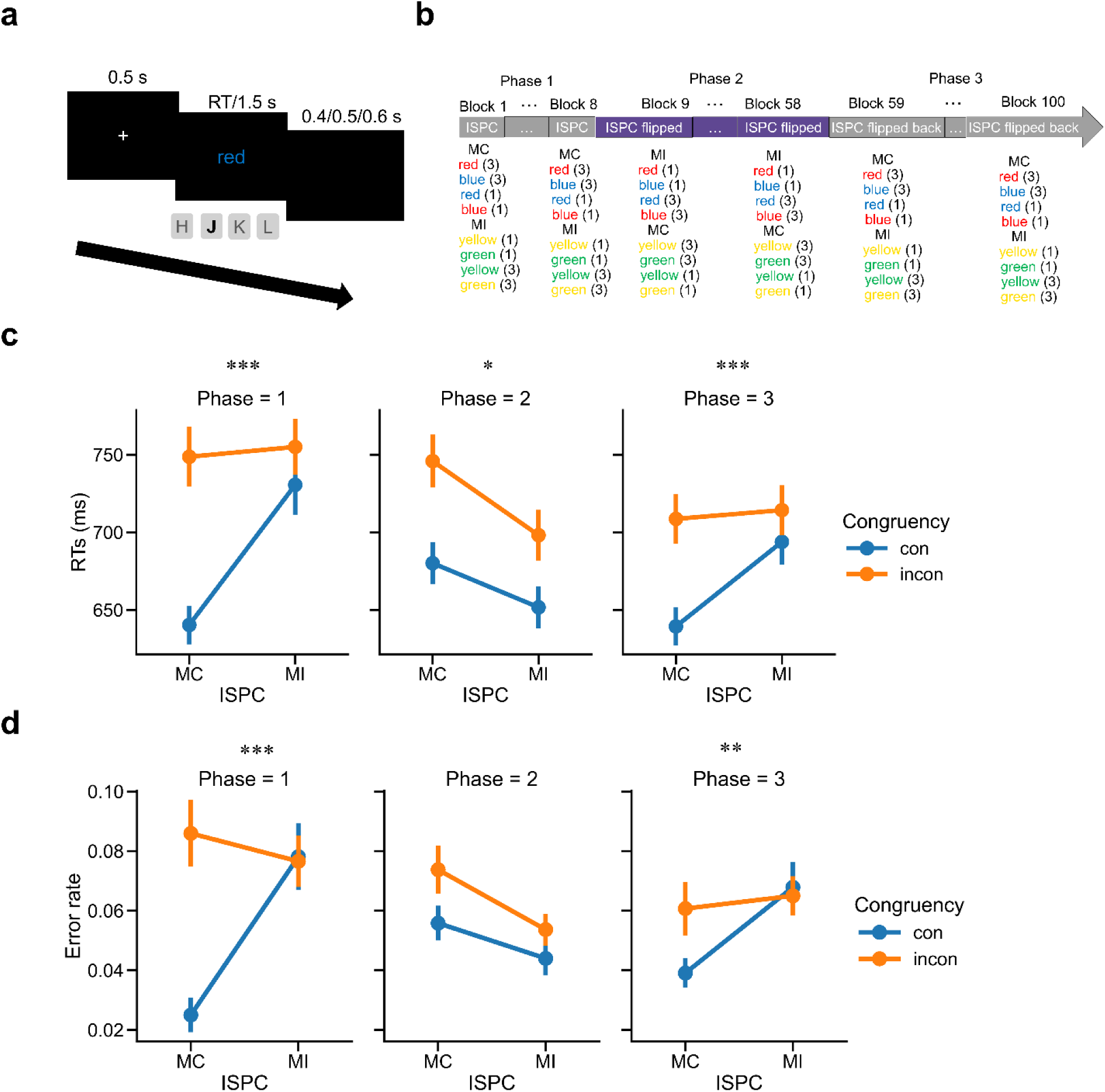
Experimental design and behavioral results (N = 40). **(a)** Trial structure. **(b)** Procedure of the main experiment. The number following each stimulus indicates its trial count within a mini-block. (**c**) Group mean RT as a function of the ISPC effect. **(d)** Group mean error rate as a function of the ISPC effect. Error bars show standard errors of the mean (SEM). MI: mostly incongruent trials; MC: mostly congruent trials; incon: incongruent trials; con: congruent trials. *: p < 0.05; **: p < 0.01; ***: p < 0.001.

### Behavioral patterns track manipulations of varying ISPC

RTs on correct trials within ±3 standard deviations (SD) of the mean and error rates were analyzed separately in each phase using a 2 (Congruency: congruent vs. incongruent) × 2 (ISPC: MC vs. MI) repeated-measures ANOVA. For RT data, there was a statistically significant interaction between Congruency and ISPC in each phase (Phase 1: *F*_(1,39)_ = 36.36, *p* < 0.001, *η*_*ρ*_^2^ = 0.482; Phase 2: *F*_(1,39)_ = 6.63, *p* = 0.014, *η*_*ρ*_^2^ = 0.145; Phase 3: *F*_(1,39)_ = 53.27, *p* < 0.001, *η*_*ρ*_^2^ = 0.577), which replicated the ISPC effect of smaller congruency effect in the MI condition compared with that in the MC condition in each phase (Phase 1: *t*_39_ = 6.03, *p* < 0.001, Cohen’s *d* = 1.174; Phase 2: *t*_39_ = 2.58, *p* = 0.014, Cohen’s *d* = 0.476; Phase 3: *t*_39_ = 7.30, *p* < 0.001, Cohen’s *d* = 1.317; Fig. 1c; Supplementary Fig. 9a). For error rate, the 2-way interaction was significant in Phase 1 (*F*_(1,39)_ = 18.05, *p* < 0.001, *η*_*ρ*_^2^ = 0.316) and Phase 3 (*F*_(1,39)_ = 10.43, *p* < 0.01, *η*_*ρ*_^2^ = 0.211) and was driven by smaller conflict effects in the MI condition than in the MC condition (Phase 1: *t*_39_ = 4.25, *p* < 0.001, Cohen’s *d* = 0.842; Phase 3: *t*_39_ = 3.23, *p* < 0.01, Cohen’s *d* = 0.655; Fig. 1d), but was not significant in Phase 2 (*F*_(1,39)_ = 1.50, *p* > 0.05; Supplementary Fig. 9b). The above findings showed that the ISPC effect flipped from Phase 1 to Phase 2, and then reversed again in Phase 3, indicating that the experimental manipulations worked as expected. Additionally, we replicated the congruency effect (i.e., worse performance on incongruent than congruent trials) in both RT and error rate data in each phase (Supplementary Note 1 and Supplementary Table 1).

### EEG analysis overview

We started by validating that the 16 experimental conditions (8 unique stimuli × MC/MI) were represented in the EEG data. Evidence of representation was provided by above-chance decoding of the experimental conditions (Fig. 2-3). We then examined whether the SC and SR associations were separable (i.e., whether SC and SR associations were different representations of equivalent information). As our results supported separable representations of SC and SR association (Fig. 4-5), we further estimated the temporal dynamics of each representation within a trial using RSA. This analysis revealed that the temporal dynamics of SC and SR association representations overlapped (Fig. 7a-b, Fig. 8a-b). To explore the potential reason behind the temporal overlap of the two representations, we investigated whether SC and SR associations were represented simultaneously as part of the task representation, independently from each other, or competitively/exclusively (e.g., on some trials only SC association was represented, while on other trials only SR association was represented). This was done by assessing the correlation between the strength of SC and SR representations across trials (Fig. 7c, Fig. 8c). Lastly, we tested how SC and SR representations facilitated performance (Fig. 9-10).

**Figure 2.**
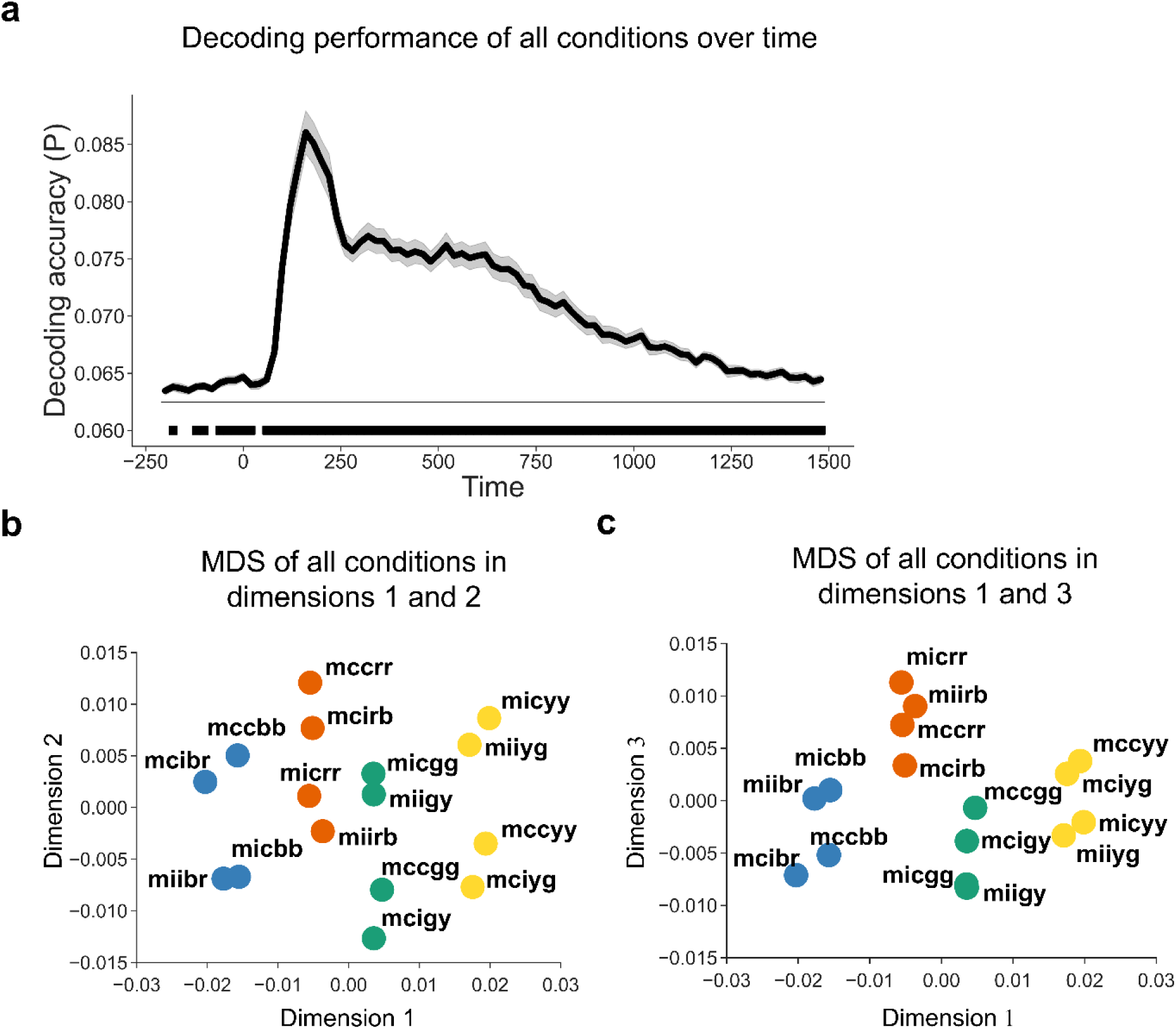
Decoding performance including both SC and SR latent subspaces (N = 40). **(a)** Group average decoding accuracy of all 16 experimental conditions as a function of time after stimulus onset. Shaded regions represent SEM. The solid line above the time axis denotes the time points showing above-chance (0.0625, or 1/16) decoding accuracy (cluster-based permutation test, cluster-forming threshold p < 0.001, cluster-level p < 0.05). **(b, c)** MDS of EEG data across all experimental conditions. The label of each dot encodes the condition in the experimental design in the order of ISPC, congruency, color, and word. For example, “mccbr” means the condition with MC, congruent trial, blue color, and the word “red”. The color of each dot denotes the ink color of each condition.

**Figure 3.**
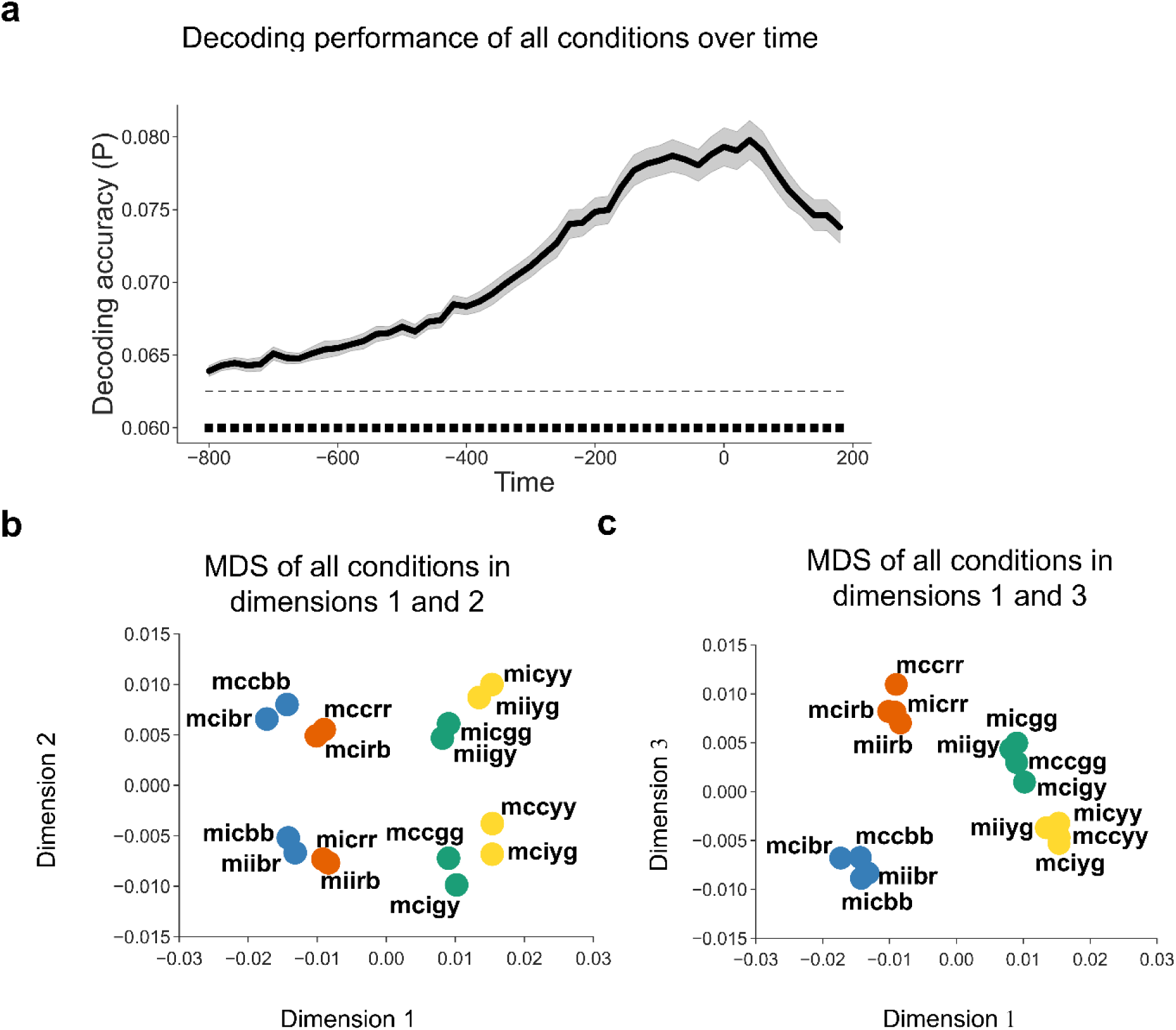
Decoding performance including both SC and SR latent subspaces using response-locked analysis (N = 40). **(a)** Group average decoding accuracy of all 16 experimental conditions as a function of time after response onset. Shaded regions represent SEM. The solid line above the time axis denotes the time points showing above-chance (0.0625, or 1/16) decoding accuracy (cluster-based permutation test, cluster-forming threshold p < 0.001, cluster-level p < 0.05). **(b, c)** MDS of EEG data across all experimental conditions. The label of each dot encodes the condition in the experimental design in the order of ISPC, congruency, color, and word. For example, “mccbr” means the condition with MC, congruent trial, blue color, and the word “red”. The color of each dot denotes the ink color of each condition.

**Figure 4.**
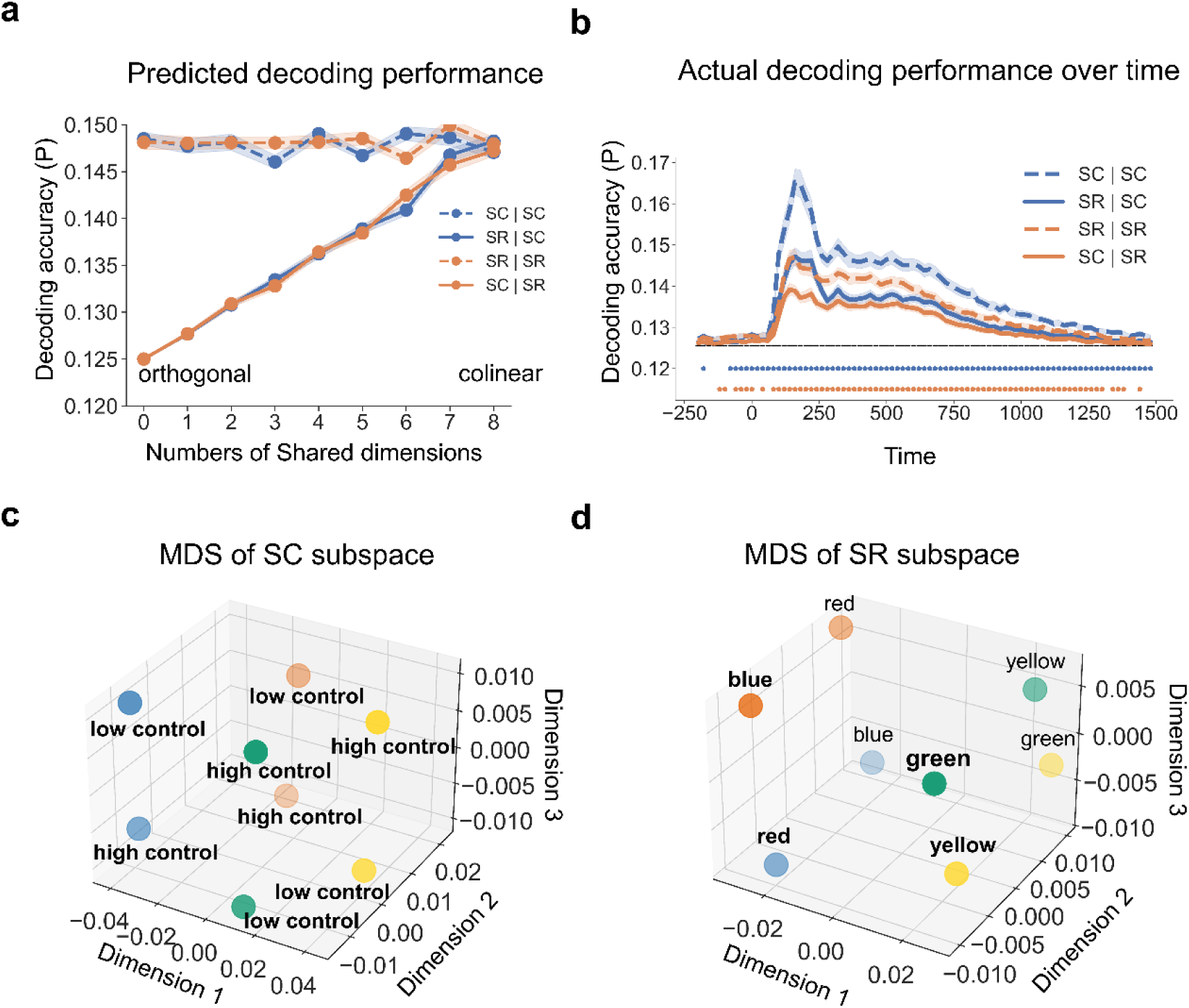
Partially overlapping SC and SR subspaces. **(a)** Simulation results (N = 40) of decoding accuracy as a function of the degree of subspace overlap, subspace type, and decoder. Both SC and SR subspaces are 8-dimensional (SC: 4 colors × MC/MI; SR: 4 words × 2 possible responses per word). The number of Shared dimensions indicates how many dimensions overlap between SC and SR subspaces. Each condition label is encoded in the format of “subspace | decoder”. For example, “SC | SR” means a SR decoder trained on the SC subspace. **(b)** Group average decoding accuracy over time as a function of which subspace the decoders are trained on. Shaded regions represent the SEM. Blue (orange) points denote the time points showing significantly better decoding accuracy when using the same subspace than when using the other subspace (cluster-based permutation test, cluster-forming threshold p < 0.001, cluster-level p < 0.05). (**c**) MDS of the SC subspace. Each dot represents the center of a SC class. Dot color and label encode the ink color and cognitive control state, respectively. **(d)** MDS of the SR subspace. Each dot represents the center of a SR class. The label and dot color encode the word meaning and associated response, respectively.

**Figure 5.**
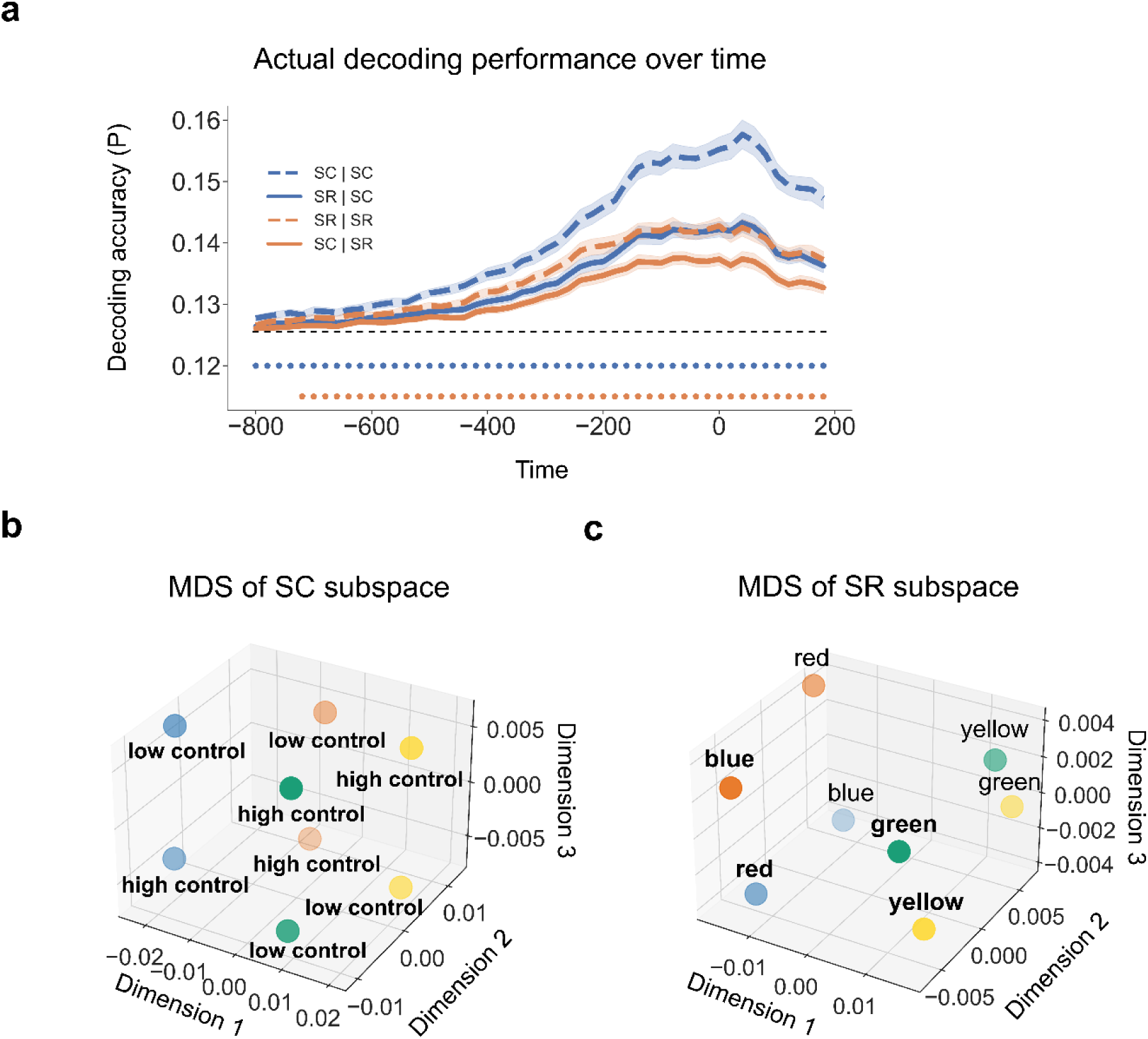
Partially overlapping SC and SR subspaces using response-locked analysis. **(a)** Group average decoding accuracy over time as a function of which subspace the decoders are trained on. Shaded regions represent the SEM. Blue (orange) points denote the time points showing significantly better decoding accuracy when using the same subspace than when using the other subspace (cluster-based permutation test, cluster-forming threshold p < 0.001, cluster-level p < 0.05). **(b)** MDS of the SC subspace. Each dot represents the center of a SC class. Dot color and label encode the ink color and cognitive control state, respectively. **(c)** MDS of the SR. Each dot represents the center of a SR class. The label and dot color encode the word meaning and associated response, respectively.

### Decodable and separate representations of controlled and non-controlled associations

Before exploring the relationship between controlled and non-controlled representations, we first confirmed that each representation was encoded during task performance. We applied linear discriminant analysis (LDA) (James et al., 2013) with the features of 64-channel event-related potentials (ERP) data to decode the 16 experimental conditions (8 unique stimuli × MC/MI) at each time point. The decoding analysis was performed on a time window from 200 ms prior to stimulus onset to 1500 ms post-stimulus onset (cf., Fig. 1a). We found that the decoding accuracy was significantly above chance level from 60 to 1500 ms after stimulus onset (Fig. 2a). To illustrate the distribution of data in the feature space, we applied multi-dimensional scaling (MDS, see Methods) to the ERP data averaged over 60 - 1500 ms after trial onset and visualized the centers of the 16 experimental conditions using the first 3 dimensions. The cumulative explained variance of the first 3 dimensions in the MDS was 0.59, 0.76 and 0.87, respectively.

Both the 8 SC classes of 16 conditions and the conditions as 4 color clusters were categorized along dimension 1 and dimension 2 (Fig. 2b) or along dimension 1 and dimension 3 (Fig. 2c). We also performed the above analyses on response-locked data and found similar patterns (Fig. 3): decoding accuracy was significantly above the chance from ∼ 800 to ∼200 ms relative to response (Fig. 3a); The cumulative explained variance of the first 3 dimensions in the MDS was 0.53, 0.70 and 0.85, respectively, revealing that MDS separated 8 SC classes of 16 conditions along dimension 1 and 2 (Fig. 3b) and categorized the conditions as 4 clusters along dimension 1 and 3 (Fig. 3c).

Prior to testing whether controlled and non-controlled associations were represented simultaneously, we first tested whether the two representations were separable in the EEG data. In other words, we reorganized the 16 experimental conditions into 8 conditions for SC (4 colors × MC/MI, while collapsing across SR levels) and SR (4 words × 2 possible responses per word, while collapsing across SC levels) associations separately. If SC and SR associations are not separable, it follows that they encode the same information, such that both SC and SR associations can be represented in the same subspace (i.e., by the same information encoded in both associations). For example, because (1) the word can be determined by the color and congruency and (2) the most-likely response can be determined by color and ISPC, the SR association (i.e., association between word and most-likely response) can in theory be represented using the same information as the SC association. On the other hand, if SC and SR associations are separable, they are expected to be represented in different subspaces (i.e., the information used to encode the two associations is different). Notably, if some, but not all, information is shared between SC and SR associations, they are still separable by the unique information encoded. In this case, the SC and SR subspaces will partially overlap but still differ in some dimensions. To summarize, whether SC and SR associations are separable is operationalized as whether the associations are represented in the same subspace of EEG data. To test this, we leveraged the subspace created by the LDA (see Methods). Briefly, to capture the subspace that best distinguishes our experimental conditions, we trained SC and SR decoders using their respective aforementioned 8 experimental conditions. We then projected the EEG data onto the decoding weights of the LDA for each of the SC and SR decoders to obtain its respective subspace. We hypothesized that if SC and SR subspaces are identical (i.e., not separable), SC/SR decoding accuracy should not differ by which subspace (SC or SR) the decoder is trained on. For example, SC decoders trained in SC subspace should show similar decoding performance as SC decoders trained in SR subspace. On the other hand, if SC and SR association representations are in different subspaces, the SC/SR subspace will not encode all information for SR/SC associations. As a result, decoding accuracy should be higher using its own subspace (e.g., decoding SC using the SC subspace) than using the other subspace (e.g., decoding SC using the SR subspace). We used cross-validation to avoid artificially higher decoding accuracy for decoders using their own subspace (see Methods). We verified this hypothesis in computational simulations to understand how the representational subspace analysis should perform under different accounts (see Methods). The simulation results (Fig. 4a) showed that: (1) when SC and SR subspaces are completely separable (i.e., the two subspaces shared no dimensions), decoding accuracy using the other subspace is at chance level; (2) when the SC and SR subspaces partially overlap (i.e., share some but not all dimensions), decoding accuracy using the other subspace is above chance level but less than when using its own subspace; and (3) when the SC and SR subspaces fully overlap (i.e., share all dimensions), decoding accuracy using the other subspace is above chance level and is comparable to that when using its own subspace.

In the EEG data (Fig. 4b; See Fig. 5a from response-locked analysis), we observed that the decoders using the other subspace (i.e., SC | SR and SR | SC) showed significantly above-chance accuracy after ∼100 ms following stimulus onset (p < 0.05, cluster-based permutation test). During the same time window, the decoder performance was significantly worse than the decoders using their own subspace (i.e., SC | SC and SR | SR). The empirical results are most consistent with the simulations of partially overlapped SC and SR spaces, suggesting that SC and SR association representations are separable in the EEG data. To show how each of the 8 conditions in the SC and SR subspaces are represented. The MDS approach was used for visualization to preserve dissimilarity between conditions when projecting from data from a high dimensional to a low dimensional space. The results showed that the 8 conditions were separate in both SC subspace (Fig. 4c, cumulative explained variance was 0.75, 0.88, 0.94 for the first 3 dimensions, respectively; See Fig. 5c from response-locked analysis, cumulative explained variance was 0.69, 0.86, 0.95 for the first 3 dimensions, respectively) and SR subspace (Fig. 4d, cumulative explained variance was 0.84, 0.93, 0.98 for the first 3 dimensions, respectively; See Fig. 5d from response-locked analysis, cumulative explained variance was 0.82, 0.95,0.99 for the first 3 dimension, respectively), which revealed the different structures of SC and SR encoded in the brain. We conducted RSA on SR decoders on the SR subspace and confirmed that these word pairs were indeed linearly separable in the SR subspace (Supplementary Fig. 1, 2).

We performed control analyses to confirm whether decoding was confounded by temporally structured noise (TSN) due to different MC and MI assignments on different phases in the design. To reduce the influence of TSN, we maximized the temporal distance between training and test data (the distance between the centers of the training and test data of the same SC/SR manipulation is about 400 trials given the experimental design) by dividing the EEG data in phase 2 and phase 1&3 into the first and second half chronologically. Phases 1 and 3 were combined because they share the same MC and MI assignments. We then trained the decoders on one half and tested them on the other half. Finally, we averaged the decoding results across all possible assignments of training and test data. This approach, like cross-phase validation in concept, revealed similar patterns (Fig. 6a-b) to the above representational subspace analyses. Additionally, the idea that TSN biases decoding results would predict a distance effect, which predicts that if a test trial is closer to a training trial with the same trial type, the higher similarity in TSN between the training and test data would more strongly inflate the decoding accuracy of the test trial, resulting in a negative correlation between distance between a test trial and its closest training trial of the same type and the test trial’s decoding accuracy. To test this predicted negative correlation, in each fold and each repetition of the cross-validation reported in the SC | SC and SR | SR decoders, we calculated the distance (mean=5.84 trials, SD=2.05, 5th percentile =2.87, 95th percentile=9.45, one trial = 2.4-2.6s) between each test trial and its closest training trial of the same trial type. This distance was used as the predictor to predict decoding accuracy in a linear regression. The regression coefficient is averaged across cross-validation folds and repetitions for each subject to match how the decoding accuracy was reported in the main text. Finally, the averaged regression coefficient was tested against 0 using a one-sample t-test. This analysis was conducted at each time point separately. As shown in the figure 6c, no time point exhibited the negative correlation as predicted by the TNS bias. In summary, our control analyses showed that the separable representations of SC and SR in the present project were not systematically biased by TSN.

**Figure 6.**
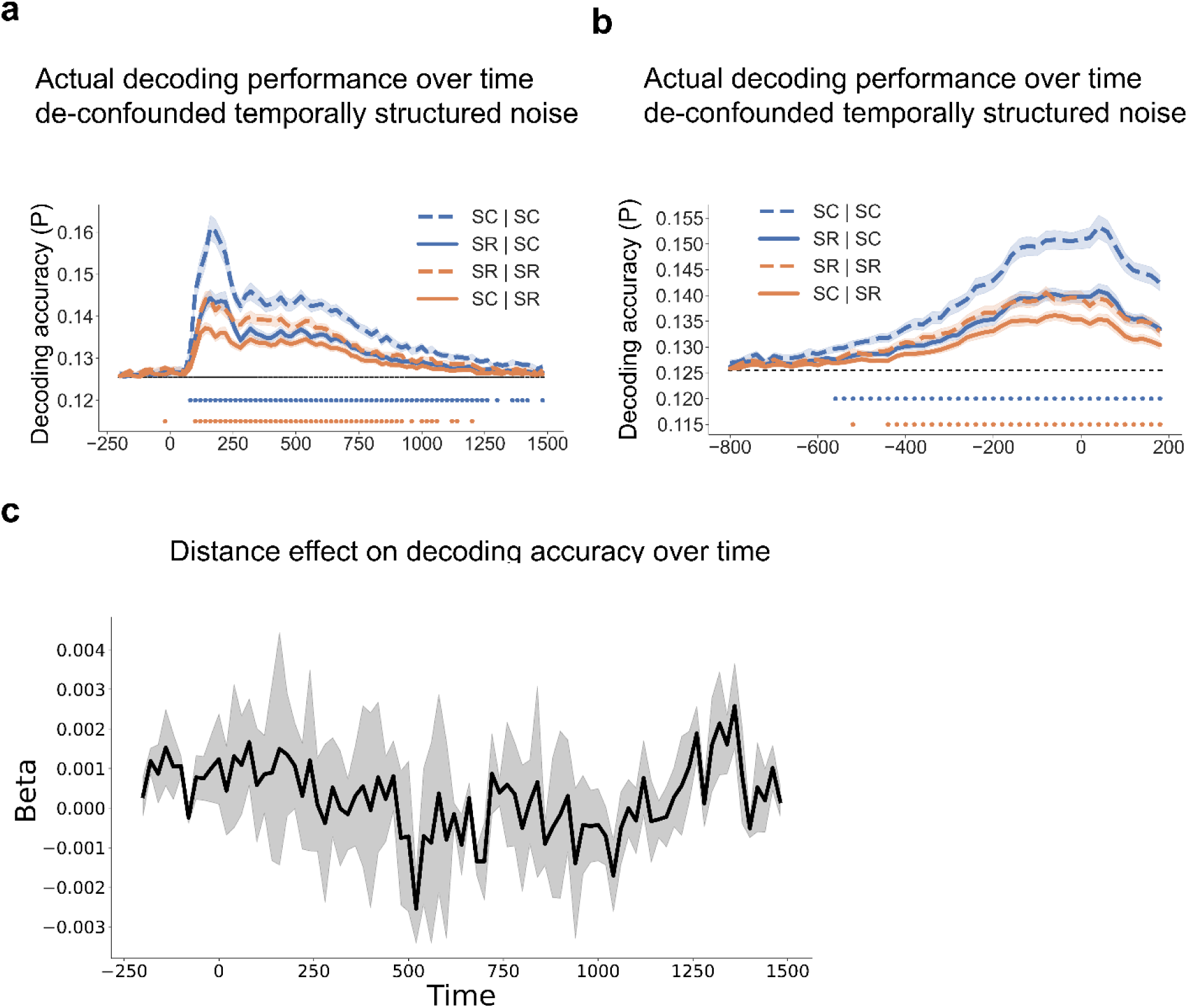
Split-half decoding and distance regression support separable SC and SR subspace. **(a, b)** Group average decoding accuracy over time is plotted as a function of which subspace the decoders are trained on (stimulus-locked data (a), response-locked data (b)). Shaded regions represent the SEM. Blue (orange) points denote the time points showing significantly better decoding accuracy when using the same subspace than when using the other subspace (cluster-based permutation test, cluster-forming threshold p < 0.001, cluster-level p < 0.05). **(c)** Slope of how much decoding accuracy changes as a function of the distance between each test trial and its closest training trial (group mean and SEM), plotted as a function of time relative to stimulus onset.

To characterize the nature of the shared features between SR and SC, we first identified which factors can be shared across SC and SR subspaces. For SC, the eight conditions are the four colors × ISPC. Thus, the possible shared dimensions are color and ISPC. Additionally, because the four colors and words are divided into two groups (e.g., red-blue and green-yellow, counterbalanced across subjects, see Methods), the group is a third potential shared dimension. Similarly, for SR decoders, potential shared dimensions are word, ISPC and group. Note that each class in SC and SR decoders has both congruent and incongruent trials. Thus, congruency is not decodable from SC/SR decoders and hence unlikely to be a shared dimension in our analysis. To test the effect of sharing for each of the potential dimensions, we performed RSA on decoding results of the SC decoder trained on SR subspace (SR | SC) (Supplementary Fig. 3a) and the SR decoder trained on SC subspace (SC | SR) (Supplementary Fig. 3b). The RSA results revealed that the contributions of group to the SC decoder (Supplementary Fig. 4a) and the SR decoder (Supplementary Fig. 4b) were significant. Meanwhile, a wider time window showed significant effect of color on the SC decoder (approximately 100 - 1100 ms post-stimulus onset, Supplementary Fig. 4a) and a narrower time window showed significant effect of word on SR decoder (approximately 100 - 500 ms post-stimulus onset, Supplementary Fig. 4b). However, we found no significant effect of ISPC on either SC or SR decoders. We also performed the same analyses on response-locked data from the time window −800 to 200 ms. The results showed shared representation of color in the SC decoder (Supplementary Fig. 4c) and group in both decoders (Supplementary Fig. 4c-d). Overall, the above results demonstrated that color, word and group information are shared between SC and SR subspaces.

In summary, our main hypothesis for the cross-subspace decoding analysis is that SR and SC subspaces are not identical. This hypothesis was supported by lower decoding accuracy for cross-subspace than within-subspace decoders and enables following analyses that treated SC and SR as separate representations.

### Simultaneously represented and positively correlated controlled and non-controlled associations

To further test whether SC and SR associations are simultaneously represented and contribute to adaptive task focus, we performed time-resolved RSA on decoding results to estimate the strength of representations of SC and SR associations while controlling for potential confounds (see Methods). In particular, since both SC and SR associations depend on the same ISPC condition, we included ISPC as a covariate in the same models as SC and SR associations to account for shared variability in the regression. The results showed that all the factors showed statistically significant effects on representational patterns (Fig. 7a). Notably, statistically significant representations of color and congruency were observed for about 1s. This long period of representation may be attributed to the trials with slow responses, as both color and congruency are crucial task information to produce the correct response and to suppress distraction from the word. Another non-exclusive possibility is that color and congruency information remained present for post-trial learning of the SC association. Crucially, both SC and SR effects remained statistically significant from ∼250 to ∼ 700 ms following the stimulus onset (Fig. 7a-b). To control for collinearity between the predictors in the RSA (see Fig. 12), we calculated the variance inflation factor (VIF) for each predictor and repeated the same analysis using predictors with VIF scores no greater than 5 (only the Identity predictor was removed). We observed similar patterns (Supplementary Fig. 5a-b), suggesting that SC and SR associations were simultaneously represented.

**Figure 7.**
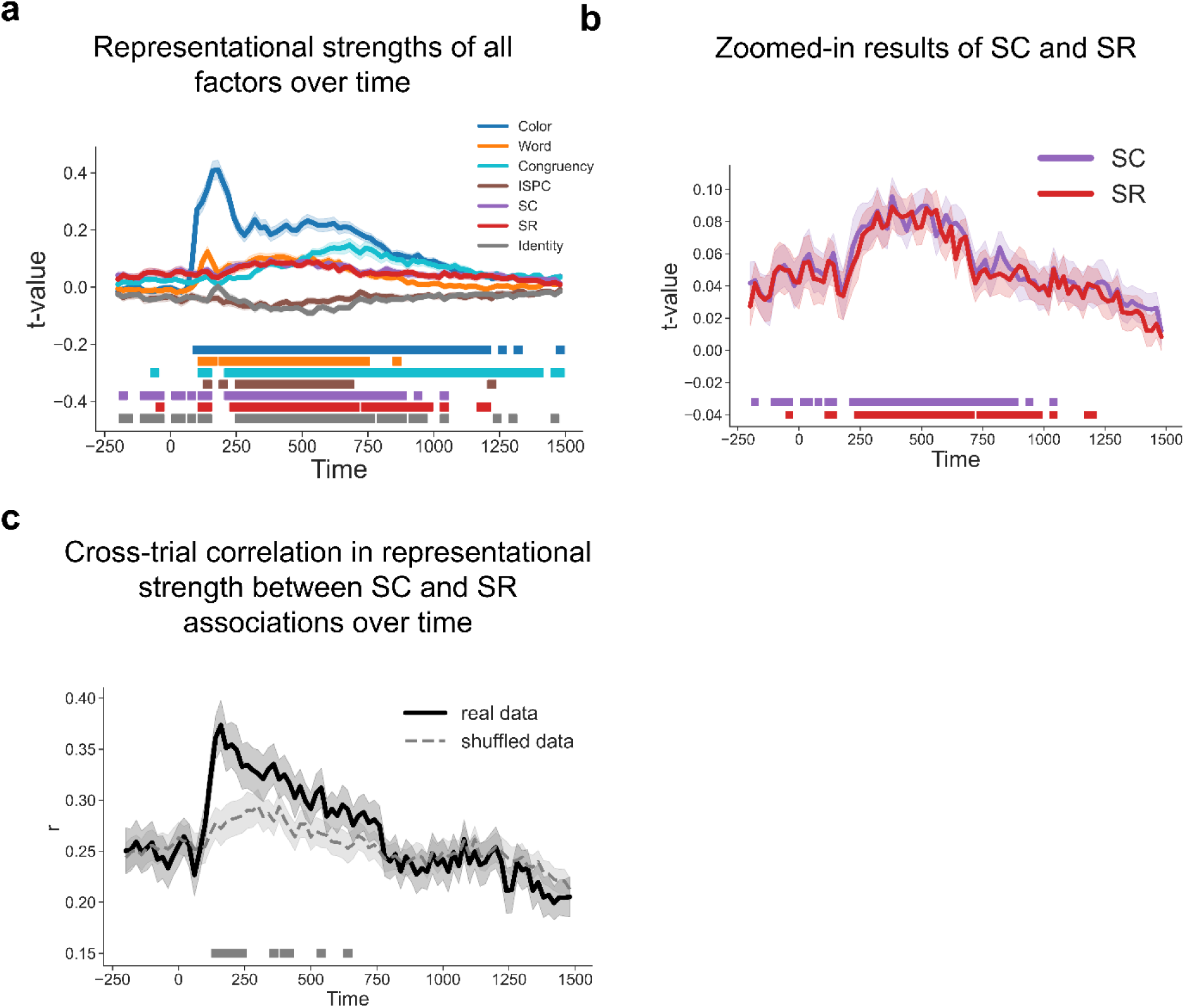
Simultaneous EEG representations of SC and SR associations. **(a)** Group average t values of representational strength for each factor over time. Squares below the lines indicate the significant time points (cluster-based permutation test, cluster-forming threshold p < 0.001, cluster-level p < 0.05). **(b)** SC and SR association results from Fig. 7a. Shaded areas denote SEM. **(c)** Cross-trial correlation coefficient of representational strength between SC and SR associations, plotted as a function of time after stimulus onset. Shaded areas indicate SEM. Squares below the lines indicate the time points significantly above the baseline correlation strength of shuffled data (Cluster-based permutation test, cluster-forming threshold p < 0.001, cluster-level p < 0.05).

The integrated task representation hypothesis predicted that the representational strength of SC and SR associations should be positively correlated across trials. Alternatively, it remained possible that at the trial level SC and SR were not represented simultaneously. That is, a trial might only show an SR effect but no SC effect, or vice versa. If this were true, SC and SR effects should be negatively correlated across trials. A third possibility is that SC and SR association representations are independent from each other, leading to no correlation. To control for potential inflation of correlation coefficients due to inherent covariance between observations (Cai et al., 2019), we randomly shuffled the trial types within each block and repeated the same analysis using the shuffled data for 10 times. The results were averaged to form a baseline correlation coefficient. We observed significantly above-baseline positive correlation between ∼100 and ∼ 450 ms following stimulus onset (Fig. 7c) and between −180 and + 50 ms relative to response (Fig. 8c). These findings indicated that trials showing a strong SC effect tended to show stronger SR effects as well, supporting the hypothesis of simultaneously encoded SC and SR associations within an integrated task representation.

**Figure 8.**
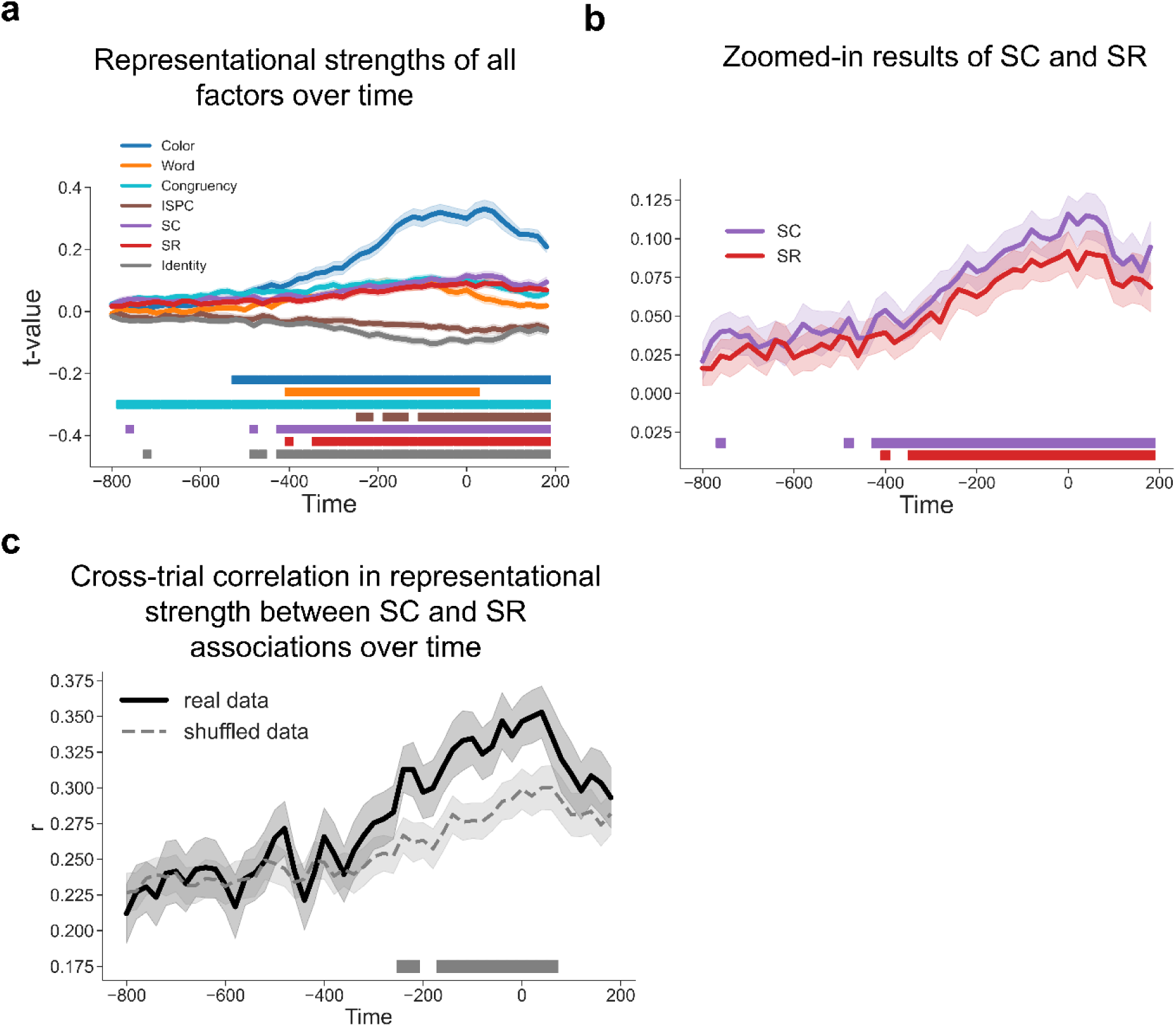
Simultaneous EEG representations of SC and SR associations using response-locked analysis. **(a)** Group average t values of representational strength for each factor over time. Squares below the lines indicate the significant time points (cluster-based permutation test, cluster-forming threshold p < 0.001, cluster-level p < 0.05). **(b)** SC and SR association results from Fig. 8a. Shaded areas denote SEM. **(c)** Cross-trial correlation coefficient of representational strength between SC and SR associations, plotted as a function of time after response onset. Shaded areas indicate SEM. Squares below the lines indicate the time points significantly above the baseline correlation strength of shuffled data (Cluster-based permutation test, cluster-forming threshold p < 0.001, cluster-level p < 0.05). The dashed line shows baseline correlation using shuffled data.

### Controlled and non-controlled associations jointly predict behavioral performance

To test whether the SC and SR effects from the RSA exhibited behavioral relevance, we predicted RT using a linear mixed model (LMM, see Methods) that included the representational strength of SC and SR associations, as quantified by the t-statistic from a trial-wise RSA. This analysis was conducted at each time point (time-locked to stimulus onset) prior to the group average RT (700.56ms) to test the effect of SC and SR representations on behavioral guidance. We found that faster responses were correlated with stronger representations of SC in the time window 380 - 700 ms (peak time at 520 ms: b = −0.045, SE = 0.006, *t* = −7.02) and SR in the time window 440 - 580 ms (peak time at 560 ms: b = −0.020, SE = 0.007, *t* = −2.93, Fig. 9a-b; See Fig. 10a-b for response-locked analysis; See Supplementary Fig. 7 and Supplementary Fig. 8 for the effects on frequent and infrequent trials). These results suggest that controlled and non-controlled associations jointly influence performance before decision-making.

**Figure 9.**
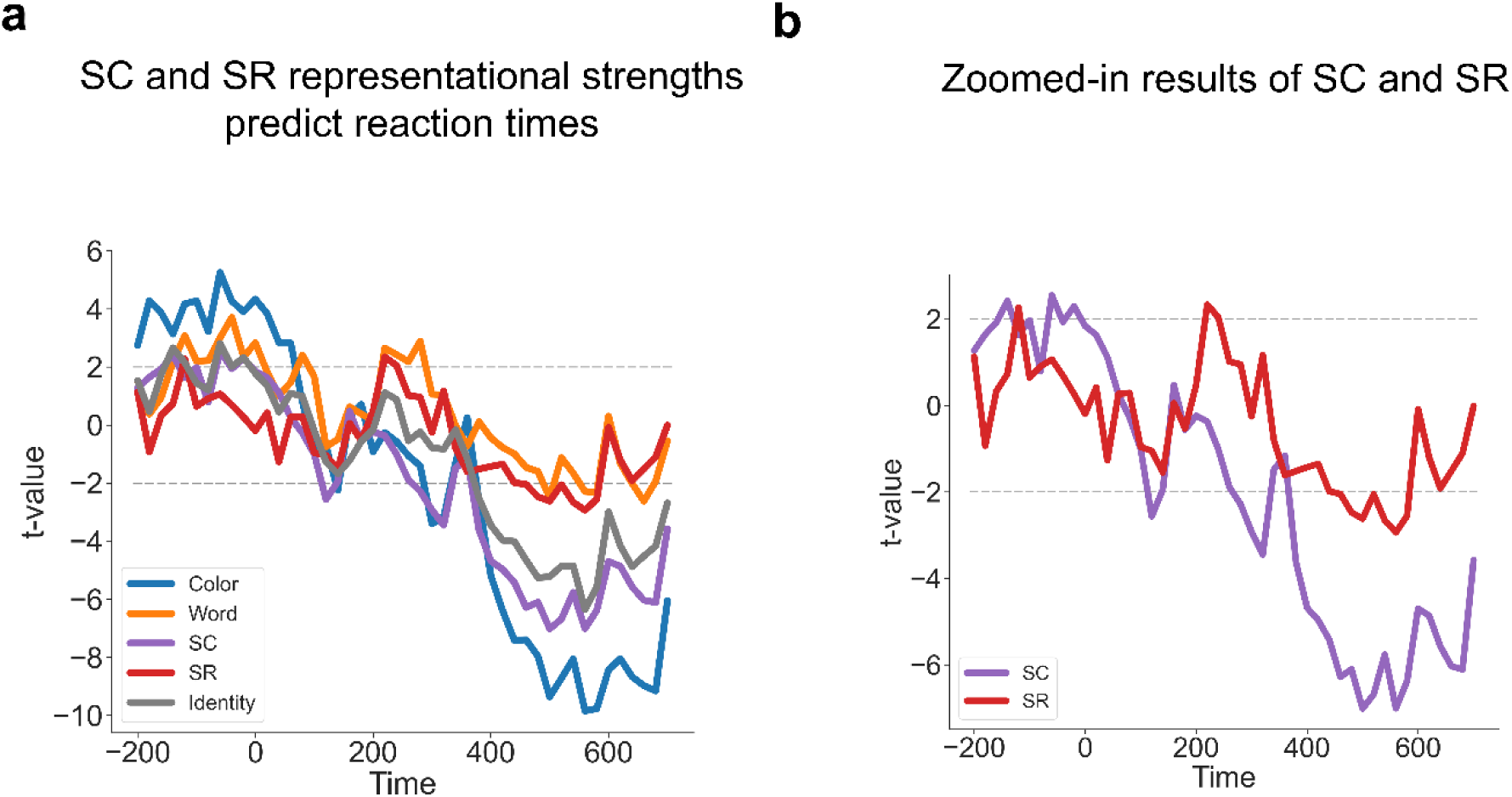
Both the strengths of SC and SR associations are correlated with RT. (**a**) Group averaged t-values for each factor predicting RT in the LMM analysis. (**b**) SC and SR association results from Fig. 9a.

**Figure 10.**
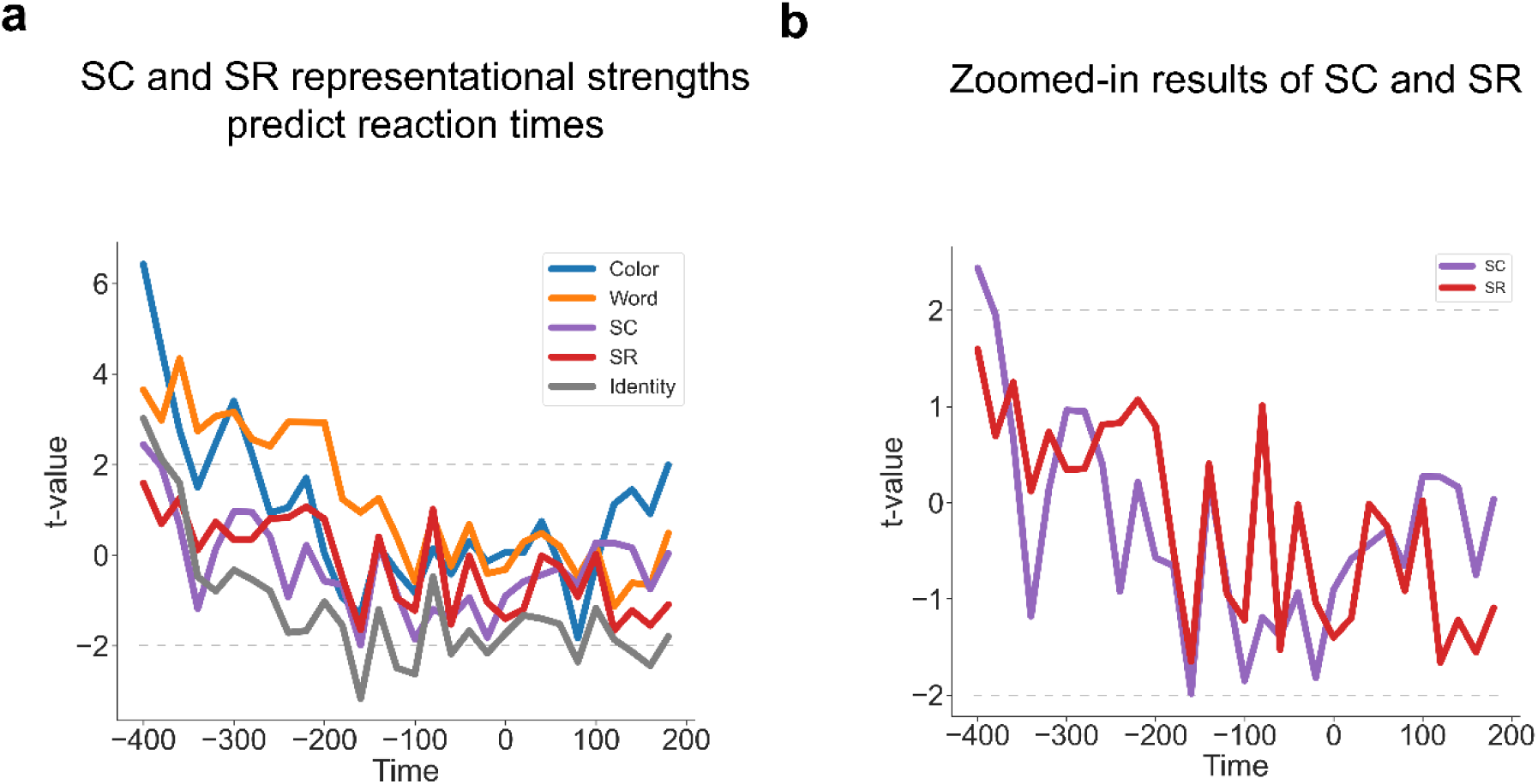
Correlations of SC and SR association strengths with RT using response-locked analysis. **(a)** Group averaged t-values for each factor predicting RT in the LMM analysis. **(b)** SC and SR association results from Fig. 10a.

## Discussion

Adaptive behavior can be guided by associations for controlled processing (e.g., stimulus-control associations) and non-controlled processing (e.g., stimulus-response associations). To explore the relationship between these representations, we combined (1) a novel extension of a classic assay of associative learning in cognitive control with (2) neural subspace analyses of the time-resolved task representations. We found consistent support for distinct neural representations of SR and SC associations, but these associations appeared to share a synchronized temporal profile consistent with them constituting a common task representation.

We found behavioral ISPC effect in each phase of our task, even after the ISPC manipulations were flipped. This suggests that participants adjusted to the local ISPC conditions, which critically allowed us to dissociate stimulus features from their SC/SR associations. The behavioral ISPC pattern was consistent with previous studies (Bejjani et al., 2020; Bugg & Hutchison, 2013; Bugg et al., 2011; Chiu et al., 2017; Ileri-Tayar et al., 2024; Ileri-Tayar et al., 2022; Jacoby et al., 2003; Spinelli et al., 2022). However, as both SC and SR associations are characterized by the same pattern of the ISPC effect, behavioral analysis and conventional contrasts between experimental conditions are insufficient to dissociate SC from SR effects. We addressed this issue with decoding, representational subspace analysis, and RSA on decoding results. We note that the strength of the ISPC effect appeared to be weaker in Phase 2 than other phases, suggesting that the neural effects of SC and SR representations may not be equivalent when the ISPC is flipped. We argue that this issue does not affect our analysis. This is because the manipulation of ISPC is binary. In other words, the decoders were trained to test whether the neural signals represent the two levels of ISPC (i.e., a higher vs. a lower level) differ systematically. The decoding analysis does not require that the neural effects of SC and SR must be numerically equivalent between phases (i.e., it is not necessary that the two levels are equidistant from the center point of SC/SR. Indeed, the decoding analysis only requires that the two levels are different). The same logic applies to the representational subspace analysis. As to the RSA, as can be seen in Fig. 12, the regressors are also binary, encoding whether two experimental conditions share the same SC/SR level without assuming equivalent neural effects.

We also found that SC and SR associations were encoded in distinct representations but appeared to be temporally synchronized as a task representation. Brain-behavior analysis prior to the mean RT supported the prediction that SC and SR can jointly guide goal-directed behaviors. Previous fMRI studies with typical ISPC paradigms showed that the ISPC effect was related to cognitive control area including the dorsolateral prefrontal cortex (dlPFC), the dorsal anterior cingulate cortex (dACC) and the parietal cortex (Blais & Bunge, 2010; Grandjean et al., 2013). Therefore, the results showing that SC and SR had distinct subspaces but emerged in the same time window should not suggest a paradox of findings or functional equivalence of SC and SR; rather, they highlight the brain’s ability to encode controlled and non-control information in parallel with partially overlapping representational subspaces to serve goal-directed behaviors. Additionally, the caudate nucleus associated with SC and the parietal cortex related with SR were observed in separate SC or SR conditions (Chiu et al., 2017). It is possible that SC and SR are supported by partially overlapping neural substrates with similar temporal dynamics. Direct evidence is needed to validate the possibility. The early onset of above-chance decoding accuracy may seem counter-intuitive. One potential confound that may cause this result is TSN. We did not find evidence that the TSN systematically biased the reported decoding results (Fig. 6). However, future research should consider orthogonalizing SC and SR classes from TSN at the source to prevent potential bias. Alternatively, because of the relatively high number of trials in each phase, it is possible that the participants (implicitly) learned the MC/MI manipulations and used the knowledge to guide proactive control, which is anticipatory and sustained (Braver, 2012; Khan et al., 2025) and then might be decoded early on or even before a trial. To better investigate this proactive control signal, we suggest that the current experimental design can be extended by adding catch trials, in which only the pre-stimulus fixation cross is presented. The proactive control hypothesis then predicts that: decoding accuracy should be above chance before the onset of the stimulus (as a replication of the present study) and following the onset of the pre-stimulus fixation on catch trials. We argue that the latter prediction is crucial because the catch trials have no stimulus, thus eliminating the confound of stimulus-elicited activities such as reactive control.

We further explored how controlled and non-controlled associations jointly affect performance via brain-behavioral analyses with LMM. In general, we observed that the strength of SC and SR association representations are both negatively correlated with trial-level variability of RT. There could be one alternative explanation that either SC or SR, instead of both, occurs on each trial. If this were true, the coefficients of SC and SR from RSA results should be negatively correlated or not correlated across trials (i.e., some trials show strong SC effects and weak SR effects, while other trials show the opposite). However, we found the opposite pattern, which indicates that there is no competition between SC and SR to be represented. One possibility leading to this positive correlation is that both associations are encapsulated in an integrated task representation (Hommel, 2004, 2019; Schumacher & Hazeltine, 2016b). Specifically, if the task representation is strongly activated on a trial, both SC and SR associations will also be strongly triggered, leading to a positive correlation between SC and SR effects across trials. Note that we found the negative prediction of the strength of SC and SR to RTs did not reach statistical significance in the response-locked analysis. It is possible that SC and SR representations have occurred before the stage of response processing, which is usually aligned with stimulus onset (Jiang et al., 2020a; Kang & Yu-Chin, 2024; Khan et al., 2025).

To interpret the ISPC effect, Bugg (2015) proposed the dual item-specific mechanisms whereby SC and SR associations learning dominate the ISPC effect under different situations, which was supported by behavioral and imaging evidence (Bugg & Hutchison, 2013; Bugg et al., 2011; Chiu et al., 2017). This means that SC associations would take over the adaptive effect when the frequency of incongruent trials is manipulated on task-relevant dimension, but SR associations would work when it is manipulated on task-irrelevant dimension. The results of the current study provide new temporal evidence for both control-driven ISPC and SR-driven ISPC and suggest that they can be represented simultaneously. Zooming in on the area of adaptive cognitive control, it has been debated whether adaptive task focus is actually involved with control or not, extending from ISPC to context/list-level PC (Brosowsky & Crump, 2016, 2021; Bugg, 2014; Bugg & Chanani, 2011; Bugg et al., 2022; Cohen-Shikora et al., 2019; Crump, 2016; Crump et al., 2017; Crump et al., 2006; Crump & Milliken, 2009; Crump et al., 2008; Crump et al., 2018; Hutchison, 2011; Schmidt, 2013b; Schmidt, 2014b; Schmidt, 2017; Schmidt et al., 2014; Spinelli & Lupker, 2021; Spinelli et al., 2019; Weidler et al., 2022) and the congruency sequence effect (Brosowsky & Crump, 2018; Schmidt, 2013a; Schmidt, 2014a; Schmidt, 2018; Schmidt & Weissman, 2016; Spinelli & Lupker, 2020b; Weissman et al., 2016). Researchers have gradually formed an associative learning view on adaptive task focus (Abrahamse et al., 2016; Egner, 2014, 2023). Egner (2014) assumed integrating different levels of abstraction about associations including SR and SC into the same learning scheme serves for adaptive behaviors. The current study supports this assumption directly with the finding of joint action of control and non-control within the same stimulus for adaptive behaviors.

To better understand the role of complex associative learning in cognitive control, we developed a new way to test how representations are integrated, and how multiple representations of task information collaborate to guide goal-directed behaviors(Badre, 2024; Musslick & Cohen, 2021). Previous research has employed partially overlapping tasks to test the conjunctive nature. For example, when there are two tasks producing different responses to the same input stimulus, encountering the stimulus will retrieve both task representations, leading to interference between the incompatible responses and impaired performance (Hubner & Druey, 2008; Koch et al., 2011; Korb et al., 2017; Rogers & Monsell, 1995). However, this test is confounded by the retrieval of episodic memory (e.g., the last experiences performing the tasks, rather than the task representations), which is also conjunctive (Altmann, 2011; Hommel, 2004, 2019). Additional evidence supporting conjunctive task representation comes from studies showing that task identity is still decodable from electroencephalographic (EEG) data when accounting for individual task features such as stimulus, rule, and response (Kikumoto et al., 2024; Kikumoto & Mayr, 2020; Kikumoto et al., 2022a; Kikumoto et al., 2022b; Rangel et al., 2023). This approach relies on shared representations of task features across multiple tasks. It remained understudied how multiple task components within the same task are influenced by the conjunctive task representation. In this study, we extend the test of conjunctive task representation in two ways. First, we examine stochastic task features that must be learned from multiple episodes of task execution to tease apart retrieval of task representation from episodic retrieval. Second, we test the prediction that conjunctive task representation will simultaneously encode multiple task features instead of locating task-unique representations.

Despite the theoretical differences between SR and SC associations, our subspace analysis revealed a pattern that is consistent with partially overlapping subspaces for SC and SR representations. SC and SR are different in their concepts and information processing, but they shared variance in their operational definitions by both depending on the same ISPC conditions. While this may have resulted in the partially overlapping representations, it may also reflect simultaneous retrieval to guide behavior. Critically, our RSA analyses revealed concurrent representations of SC and SR associations even when controlling for ISPC confounds. The experimental design combined with the RSA method ensures that the predictors for SC and SR associations are different.

An SR association itself is automatic behavior, but it is sometimes included as a core component of cognitive control (Badre, 2024). We consider SC and SR as controlled and uncontrolled respectively based on the literature investigating the mechanism of ISPC effect. The SC account posits that the ISPC effect results from conflict and involves conflict adaptation, which requires the regulation of attention or control (Bugg & Hutchison, 2013; Bugg et al., 2011; Schmidt, 2018; Schmidt & Besner, 2008). On the other hand, the SR account argues that ISPC effect does not require conflict adaptation but instead reflects contingency leaning. That is, the response can be directly retrieved from the association between the stimulus and the most-likely response without top-down regulation of attention or control. As more empirical evidence emerged, researchers advocating control view began to acknowledge the role of associative learning in cognitive control regarding the ISPC effect (Abrahamse et al., 2016). SC association has been thought to include both automatic that is fast and resource saving and controlled processes that is flexible and generalizable (Chiu, 2019). Overall, we do not intend to claim that SC is entirely controlled or SR is completely automatic. We use SC-controlled and SR-uncontrolled representations to align with the original theoretical motivation and to highlight the conceptual difference between SC and SR associations.

A limitation of the current design is that in theory temporally structured noise (e.g., autocorrelation in EEG data) may bias the decoding accuracy due to the blocked design. Although the present data provided no evidence that the decoding results in this study were biased by the temporally structured noise, future studies should aim to develop experimental designs that eliminate this potential confound at the source. One potential solution would be to introduce additional phases flipping ISPC manipulations. At the same time, enough trials must be included in each phase to ensure the strength of the ISPC effect within each phase. A careful balance between session number and length will be helpful to optimize the duration of such a design.

In summary, we find initial evidence of neural representations of associations of both controlled and non-controlled information quantified by SC and SR. Our results suggested that the neural representations are represented simultaneously in an integrated representation in partially overlapping neural subspaces shortly after stimulus onset and remain active after the response is made. Moreover, the strengths of the associations prior to the response seem to jointly predict performance. These results shed light on how an integrated task representation, which consists of multiple associations that can drive behavior, can retrieve multiple associations of controlled and non-controlled information to guide adaptive behaviors.

## Acknowledgements

We thank Atsushi Kikumoto and the members of the Cognitive Control Collaborative at the University of Iowa for helpful comments and suggestions. We also thank Jordan Nicholson, Tommy Looi, Yong-Yao Cheng and Qiutong Hong for their help with data acquisition. This project was supported by the National Institute of Mental Health (R01MH131559 to J.J.), and the C.V. Starr Fellowship (H.R).

## Methods

### Participants

Sixty-one healthy adults participated in the EEG experiment. Sixteen participants with excessive muscle artifacts (6 subjects), too many corrupted EEG channels (5 subjects), and missed partial data (5 subjects), and five participants with accuracy below 75% were removed, resulting in a final sample of 40 participants (28 females, 12 males; age mean = 23.9, SD = 6.2, age information was not acquired for the first 25 participants). All participants had normal or corrected-to-normal vision. All participants gave informed consent prior to the experiment, and either were paid or received course credits for participation. All procedures were approved by the University of Iowa Institutional Review Board (UIIRB #202001345).

#### Stimuli

The experiment was programmed using PsychoPy (ver. 2022.2.5). The stimulus was a color word presented in the center of the screen (24 inch LCD display with resolution of 1920× 1080) on a black background. Four color words (red, blue, yellow, and green) and their corresponding colors were used. For each participant, the colors and their corresponding words were divided into two sets of red-blue and yellow-green. Within each set, the words and colors formed a 2 × 2 factorial design, resulting in eight unique Stroop stimuli (Stroop, 1935) in total (e.g., red in red or blue color, blue in red or blue color, yellow in yellow or green color and green in yellow or green color).

### Design and procedures

Each trial started with the presentation of a fixation cross for 500 ms, followed by a color word at the center of the screen (specified in ‘height’ with 0.08 units where the full screen height equaled 1.0 unit) for 1500 ms or until a response was made (Fig. 1a). The four colors were mapped onto four keys (“H” for red, “J” for blue, “K” for yellow and “L” for green). Trials were separated by a black screen for the remainder of the 1500 ms response deadline plus a jittered (400/500/600 ms with equal probability) interval. The trials were grouped into mini-blocks of 16 trials each. The participants first underwent a practice session with 3 mini-blocks of 50% congruent trials. During practice, participants received feedback for a correct, incorrect or too slow response on each trial. After reaching the 75% accuracy criteria across all blocks, they moved on to the main experiment (Fig. 1b) with 8 mini-blocks for phase 1, 50 mini-blocks for phase 2 and 48 mini-blocks for phase 3.

### Behavioral analysis

Behavioral data from the main experiment were analyzed. For response time (RT) data, only correct trials within ±3 SD from each subject’s mean RT were included in the analysis. Accuracy and RT were analyzed separately for each phase using a 2 (congruency) × 2 (ISPC) repeated-measures ANOVA.

### EEG data acquisition and preprocessing

EEG data were recorded from a 64-channel active electrode cap with the ground at Fz and the reference at Pz using an actiCHamp amplifier (Brain Products) at a sampling rate of 500Hz with a hardware band-pass filter from 0.016 Hz to 1000 Hz. All electrode impedances werekept below 10 kΩ during the experiment. All EEG data were analyzed using the EEGLAB toolbox (https://sccn.ucsd.edu/eeglab/index.php) and custom MATLAB scripts. Preprocessing started with rejection and interpolation of noisy electrodes. Next, a 0.1–50 Hz band-pass filter was applied to the raw EEG data. The data were then re-referenced to the average of all electrodes and segmented into epochs spanning from −500 ms to 1500 ms relative to the stimulus onset. We used independent component analysis (ICA) to remove eye-blink and muscle artifacts from the epochs. These epochs were further segmented into smaller epochs ranging from −200 to 1500 ms relative to stimulus onset. These smaller epochs were also used to perform the response-locked analysis ranging from −800 to 200 ms relative to response. Epochs with amplitudes exceeding ± 80 μV were rejected. Fewer than 70% of all the epochs were excluded for each subject.

### Decoding analyses

Decoding analysis was conducted on epoched EEG data without baseline correction to avoid potential bias in the decoding results (van Driel et al., 2021). To generate the dependent variable for the RSA, we first decoded the representations of all conditions using LDA (scikit-learn version 1.2.2 in Python (Pedregosa et al., 2011)). The preprocessed 64-channel ERP data were down-sampled to 50 Hz and were used as the features to decode the 16 experimental conditions (8 unique stimuli × MC/MI. Note that Phases1 and 3 shared the same ISPC manipulations). Decoding results were obtained by performing a 4-fold cross-validation procedure (Fig. 11). To avoid the bias of unbalanced trial numbers among different classes on decoding results, the more frequent trial types were down-sampled to be consistent with the less frequent trial types. Specifically, given the 75%/25% proportion congruency used in the manipulation of ISPC, the more frequent trial types had three times as many trials as the less frequent ones. Thus, the more frequent trial types were randomly divided into three parts, and one part was used for the 4-fold cross-validation. The cross-validation was repeated 30 times (i.e., 10 random partitions of the frequent trial types and each part being used once within each random partition). The LDA decoding accuracy was averaged. The decoding analysis was performed at each time point of a trial for each participant.

**Figure 11.**
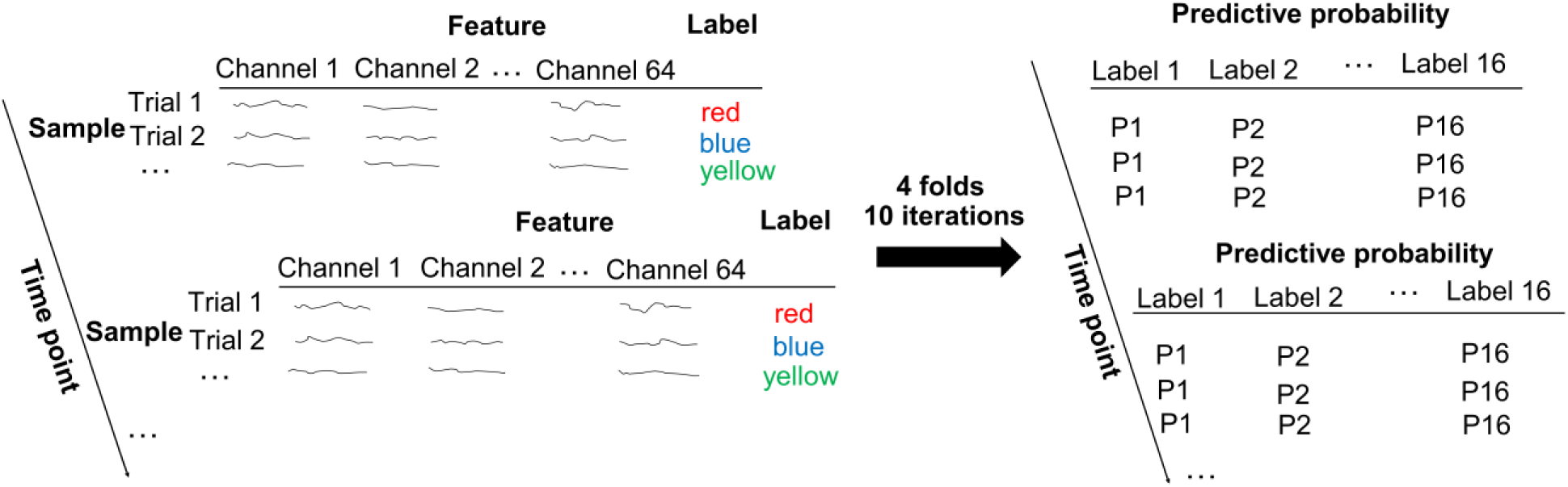
Illustration of linear discriminatory analysis (LDA).

To construct SC and SR subspaces, we trained 8-way SC (4 colors × MC/MI) and SR (4 words × 2 possible responses per word) LDA decoders and then transformed all training and test data into the subspaces of the LDA decoders. For each subspace, we built and tested SC and SR decoders using the same cross-validation approach as above. To avoid bias, all decoders used the same partition of training and test data, with the class label of each trial determined by its experimental condition. To test the statistical significance of decoding results, cluster-based permutation test (Maris & Oostenveld, 2007) with 1000 permutations was applied to address the issue of multiple comparisons. Clusters were formed with a threshold of p < 0.001 and cluster significance was assessed with a threshold of p < 0.05 (Collins & Frank, 2018). Additionally, classical multidimensional scaling (MDS) analysis (Wickelmaier, 2003) on a 16×16 matrix of probabilistic classification results from the 16-way decoders was applied to visualize the neural representations of all conditions potentially including both SC and SR associations. Similar MDS analyses were performed on the 8×8 matrix of probabilistic classification results from the SC decoder and the other 8×8 matrix of decoding accuracy from the SR decoder to visualize their neural representations.

### Simulations of cross-decoding accuracy

To simulate SC and SR subspaces with different levels of overlap, we first constructed non-overlapping SC and SR subspaces, each having 8 dimensions and consisting of 8 classes of 100 data points each. The centers of the classes were equidistant, with the distance set to the mean between-center distance in the EEG subspace. For each class, the data points were randomly drawn from a multivariate Gaussian distribution using the class center as the mean and the within-class standard deviation from the EEG subspace (averaged across all time points). Both SC and SR decoding were then conducted on each of the SC and SR subspaces using 4-fold cross-validation. To examine how overlap in subspaces affects decoding accuracy, the overlap between the two subspaces was manipulated by the number of shared dimensions. That is, if n (n ≤ 8) dimensions were shared, n dimensions of data were randomly chosen for SC_SR (i.e., SR decoder trained on the SC subspace) and SR_SC decoders. For SC_SC and SR_SR decoders, all 8 dimensions were used regardless of n.

### RSA on decoding results

To further track the representational dynamics of SC and SR associations, we tested the hypothetical representational similarity patterns using the empirical representational similarity patterns from EEG data and linear regression-based RSA (Kikumoto & Mayr, 2020; Kikumoto et al., 2022a; Rangel et al., 2023). For each time point, its decoding results reflected the neural similarity between conditions (i.e., if two conditions have similar neural representations, the misclassification rate would increase) and were organized as the logit-transformed probability of a trial belonging to each of the 16 conditions to be the dependent variable. Six factors were considered, namely color, word, congruency, ISPC (MC/MI), SC (i.e., the pairing between color and ISPC), and SR (i.e., the pairing between word and ISPC), resulting in six hypothetical representational similarity patterns in the form of binary linear regressors (Fig. 12). Specifically, for each trial, each regressor was a column-vector of 16 cells, with each cell containing a binary value indicating whether each of the 16 conditions shared the feature (e.g., color) with the condition to which the current trial belonged to. Note that, as the SR association binds the word with the most-likely response rather than the actual response on the current trial, two trial types with different responses in the RDM (Fig. 12) may share the same SR association, as long as they share the same word and the same ISPC manipulation, which will produce the same most-likely response. Additionally, two nuisance binary regressors marking the condition and its frequency (i.e., whether a MC color on a congruent trial or a MI color on an incongruent trial) of the current trial were included. For each participant, the within-trial regressors formed a design matrix to regress against the empirical decoding results, resulting in trial-wise *t* values for each regressor. For each time point, individual mean *t* values for each regressor were submitted to a group-level one-sample *t*-test against 0. To test the significance of temporal representational strength, cluster-based permutation test (1000 permutations using identical cluster forming thresholds to the main analysis) was applied to address the issue of multiple comparisons. Additionally, we correlated those regression coefficients of SC with that of SR across trials at each time point to test the possibility that SC and SR were not simultaneously represented within a trial.

**Figure 12.**
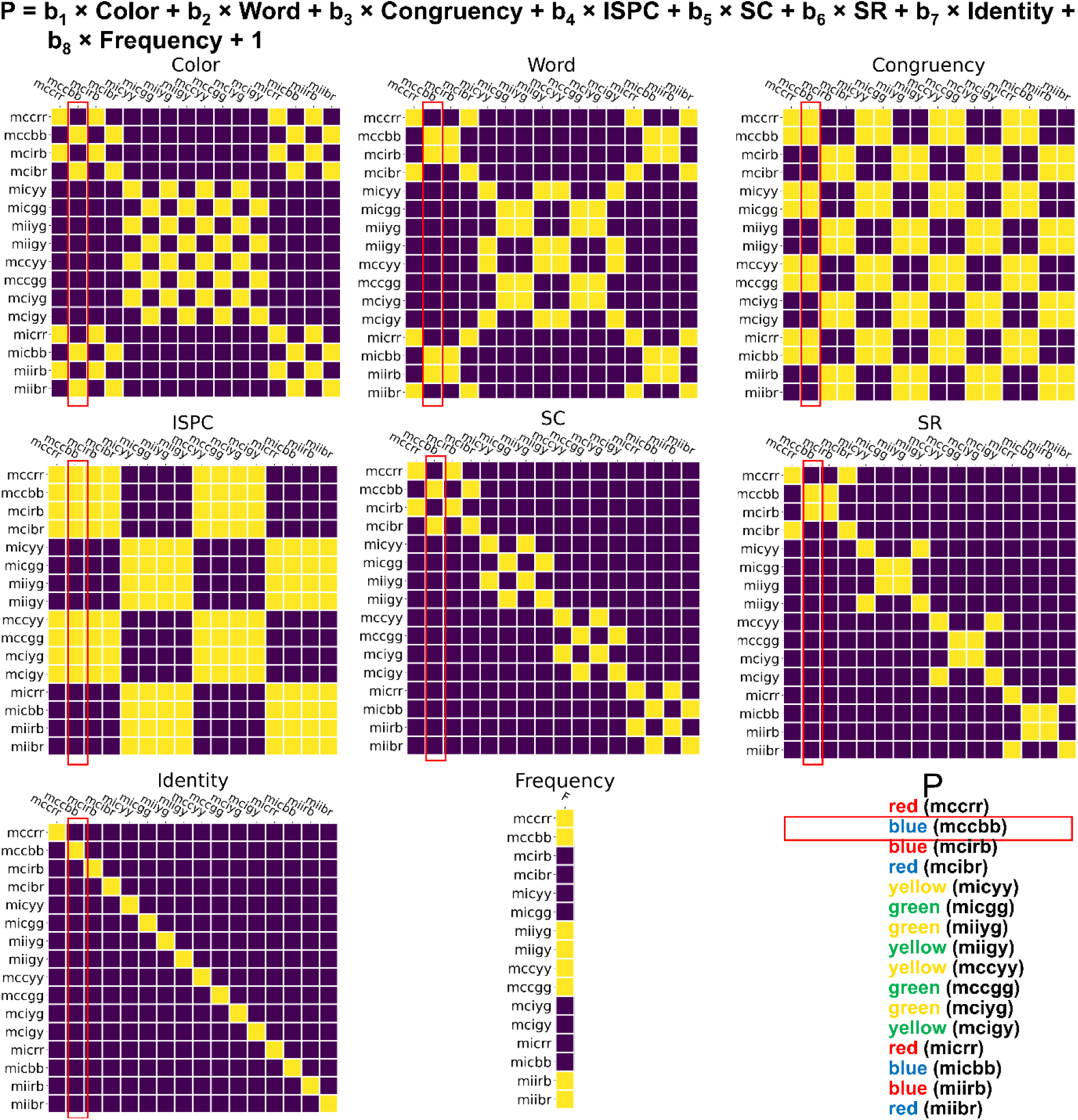
Illustration of RSA. The label of each row/column represents the condition in the experimental design including ISPC, congruency, color and word. For example, “mccbb” means the condition with MC, congruent trial, color blue and word blue. For each cell in a matrix, the color indicates whether the row and column conditions share the same factor (yellow = yes, blue = no) encoded by the matrix. For example, the cell at the 4^th^ row and the 2^nd^ column in the “SC” matrix encodes that the 4^th^ condition (i.e., mcibr) and the 2^nd^ condition (i.e., mccbb) share the same SC association.

### Linear mixed model (LMM)

We further performed LMM at each time point preceding the mean RT across all trials (700.56 ms) to test whether trial-wise RSA results could predict the RT variability (Kikumoto & Mayr, 2020; Rangel et al., 2023). The model predicted trial-wise RT using each of the eight trial types in SC or SR (t-values from the RSA regressors) as separate predictors with their own intercepts, while including random intercepts for subjects. Log-transformed RTs were used as the dependent variable. At each time point, we only included trials that have not generated a response (for stimulus-locked analysis) or already started (for response-locked analysis).

## Notes

### Competing Interest Statement

The authors have declared no competing interest.

### Summary of Updates

The manuscript was revised based on reviewers' new comments.

