## Supplementary for "Adaptive behavior is guided by integrated representations of controlled and non-controlled information"

### Supplementary materials

#### Supplementary results.

#### Behavior

| | | Phase 1 | | Phase 2 | | Phase 3 | |
|---|---|---|---|---|---|---|---|
| | | MC | MI | MC | MI | MC | MI |
| RTs (ms) | con | 640.38 (10.98) | 730.58 (18.88) | 680.15 (12.20) | 651.75 (12.10) | 639.45 (10.96) | 693.87 (13.30) |
| | incon | 748.65 (17.84) | 754.98 (16.49) | 745.86 (15.59) | 698.17 (14.87) | 708.65 (14.54) | 714.27 (14.62) |
| Error rate | con | 0.02 (0.005) | 0.08 (0.010) | 0.06 (0.005) | 0.04 (0.005) | 0.04 (0.004) | 0.07 (0.008) |
| | incon | 0.09 (0.010) | 0.08 (0.008) | 0.07 (0.007) | 0.05 (0.005) | 0.06 (0.008) | 0.06 (0.006) |

Supplementary Table 1. Descriptive statistics of behavioral results. Data are reported in the format of group mean (SEM). MI: mostly incongruent trials; MC: mostly congruent trials; incon: incongruent trials; con: congruent trials.

Note 1.

In RT data, the main effect of ISPC in each phase was significant (Phase 1: $F_{(1,39)} = 29.40$, $p < 0.001$, $\eta_\rho^2 = 0.430$; Phase 2: $F_{(1,39)} = 22.15$, $p < 0.001$, $\eta_\rho^2 = 0.362$; Phase 3: $F_{(1,39)} = 18.98$, $p < 0.001$, $\eta_\rho^2 = 0.327$), reflected by faster responses in the MC condition than in the MI condition in phases 1 and 3 (Phase 1: $t_{39} = -5.42$, $p < 0.001$, Cohen's $d = 0.508$; Phase 3: $t_{39} = -4.36$, $p < 0.001$, Cohen's $d = 0.362$) but the opposite pattern in phase 2 ($t_{39} = 4.71$, $p < 0.001$, Cohen's $d = 0.449$). The expected main effect of Congruency in each phase was also observed (Phase 1: $F_{(1,39)} = 55.56$, $p < 0.001$, $\eta_\rho^2 = 0.588$; Phase 2: $F_{(1,39)} = 115.95$, $p < 0.001$, $\eta_\rho^2 = 0.748$; Phase 3: $F_{(1,39)} = 86.71$, $p < 0.001$, $\eta_\rho^2 = 0.690$), driven by slower responses on incongruent trials than congruent trials (Phase 1: $t_{39} = 7.45$, $p < 0.001$, Cohen's $d = 0.698$; Phase 2: $t_{39} = 10.77$, $p < 0.001$, Cohen's $d = 0.680$; Phase 3: $t_{39} = 9.31$, $p < 0.001$, Cohen's $d = 0.550$).

For error rate, there were significant main effects of ISPC (Phase 1: $F_{(1,39)} = 7.60$, $p < 0.01$, $\eta_\rho^2 = 0.163$; Phase 2: $F_{(1,39)} = 11.69$, $p < 0.01$, $\eta_\rho^2 = 0.231$; Phase 3: $F_{(1,39)} = 9.54$, $p < 0.01$, $\eta_\rho^2 = 0.197$) reflected by less error-prone in the MC condition than in the MI condition in phases 1 and 3 (Phase 1: $t_{39} = -2.76$, $p < 0.01$, Cohen's $d = 0.536$; Phase 3: $t_{39} = -3.09$, $p < 0.01$, Cohen's $d = 0.437$) but reversed in phase 2 ($t_{39} = 3.42$, $p < 0.01$, Cohen's $d = 0.515$), and Congruency (Phase 1: $F_{(1,39)} = 10.55$, $p < 0.01$, $\eta_\rho^2 = 0.213$; Phase 2: $F_{(1,39)} = 10.99$, $p < 0.01$, $\eta_\rho^2 = 0.220$; Phase 3: $F_{(1,39)} = 4.26$, $p < 0.05$, $\eta_\rho^2 = 0.099$) reflected by more error-prone on incongruent trials than congruent trials (Phase 1: $t_{39} = 3.25$, $p < 0.01$, Cohen's $d = 0.687$; Phase 2: $t_{39} = 3.32$, $p < 0.01$, Cohen's $d = 0.456$; Phase 3: $t_{39} = 2.06$, $p < 0.05$, Cohen's $d = 0.255$).

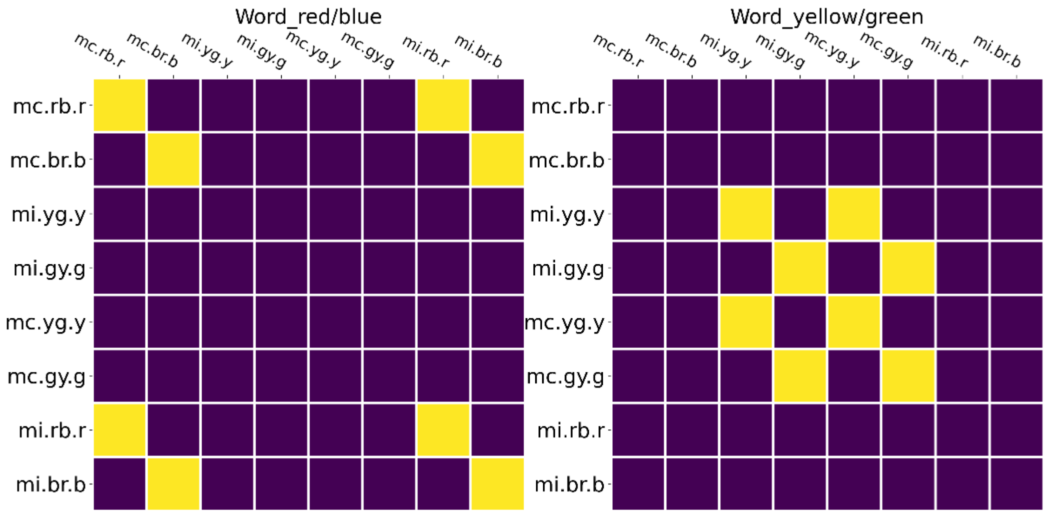


**Supplementary Figure 1 | Illustration of similarity matrices of RSA on SR decoder trained on SR subspace.** To test whether word can be linearly separated in SR subspace, we performed the RSA with word on the SR decoder trained on the SR subspace. Note that there are two pairs of words that are not linearly separable in Fig. 4d: red-blue and yellow-green. Thus, we specifically tested the separability within the two pairs using one predictor for each pair. The label of each row/column represents the condition including ISPC, color and word. For example, "mc.rb.r" means the condition with MC, color red & blue, and word red. For each cell in a matrix, the color indicates whether the row and column conditions share the same factor (yellow = yes, purple = no) encoded by the matrix. For example, the cell at the 7$^{th}$ row and the 1$^{st}$ column in the "word_red/blue" matrix encodes that the 7$^{th}$ condition (i.e., mi.rb.r) and the 1$^{st}$ condition (i.e., mc.rb.r) share the same word association.

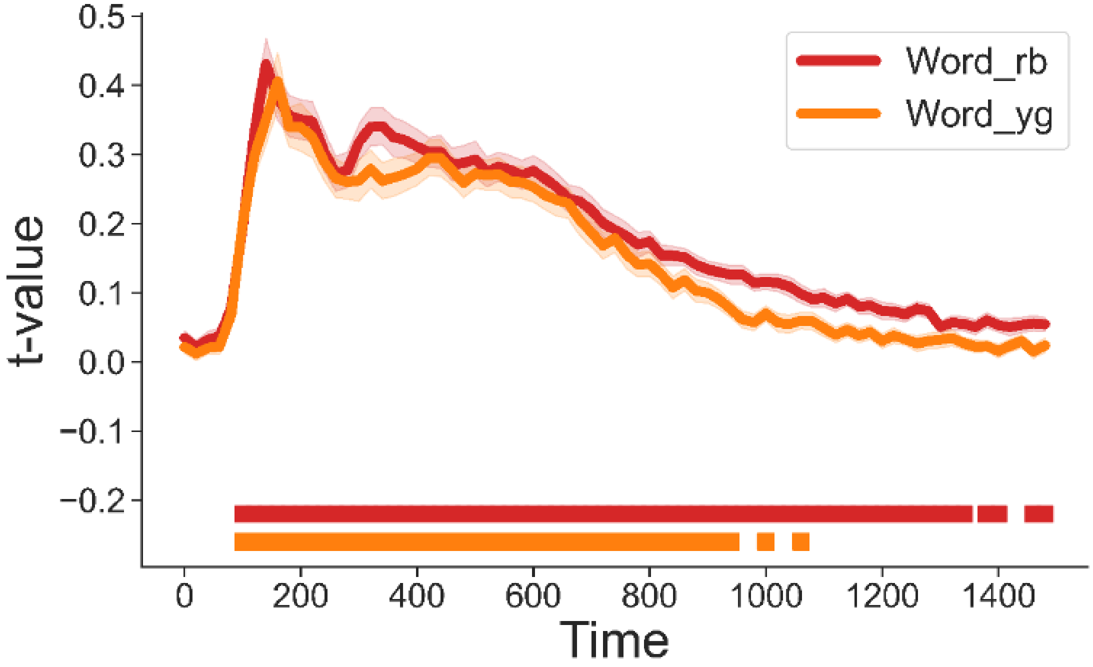


**Supplementary Figure 2 | linear separation of the word feature on SR subspace.** Group average t values of representational strength for word red & blue and word yellow & green on SR decoder trained on SR subspace. Squares below the lines indicate statistically significant time points ($p < 0.001$, Bonferroni corrected). The results showed that within both word pairs (blue-red, yellow-green) individual words were represented above chance level. Considering that the decoders were linear, this finding indicated linear separability of the word pairs in the original SR subspace.

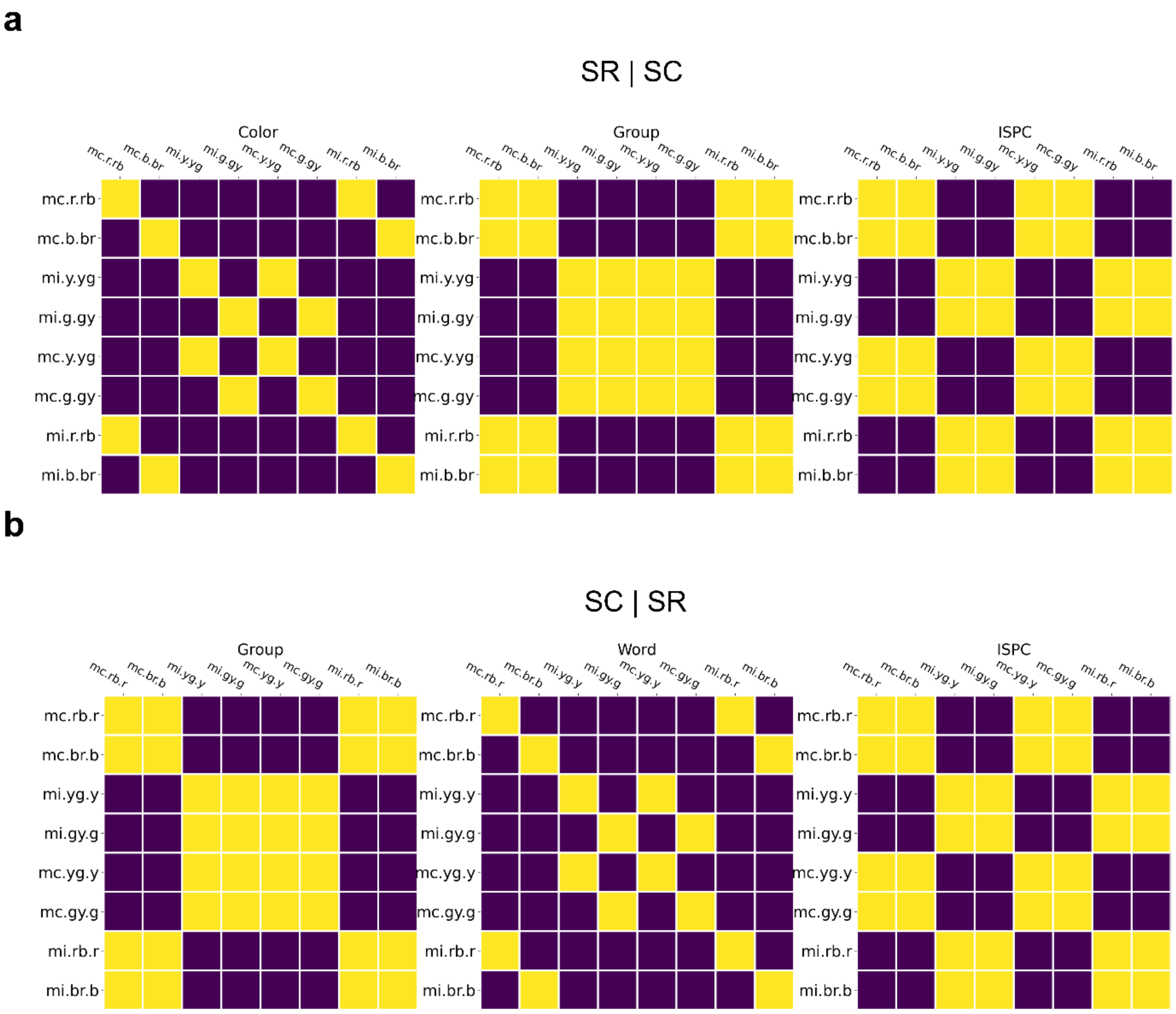


**Supplementary Figure 3 | Illustration of Similarity matrix of RSA.** (**a**) Similarity matrices of SC decoder trained on SR subspace. The label of each row/column represents the condition including ISPC, color and word. For example, "mc.r.rb" means the condition with MC, color red, and word red & blue. For each cell in a matrix, the color indicates whether the row and column conditions share the same factor (yellow = yes, purple = no) encoded by the matrix. For example, the cell at the 7$^{th}$ row and the 1$^{st}$ column in the "Color" matrix encodes that the 7$^{th}$ condition (i.e., mi.r.rb) and the 1$^{st}$ condition (i.e., mc.r.rb) share the same color association. (**b**) Similarity matrices of SR decoder trained on SC subspace. The label of each row/column represents the condition including ISPC, color and word. For example, "mc.rb.r" means the condition with MC, color red & blue, and word red.

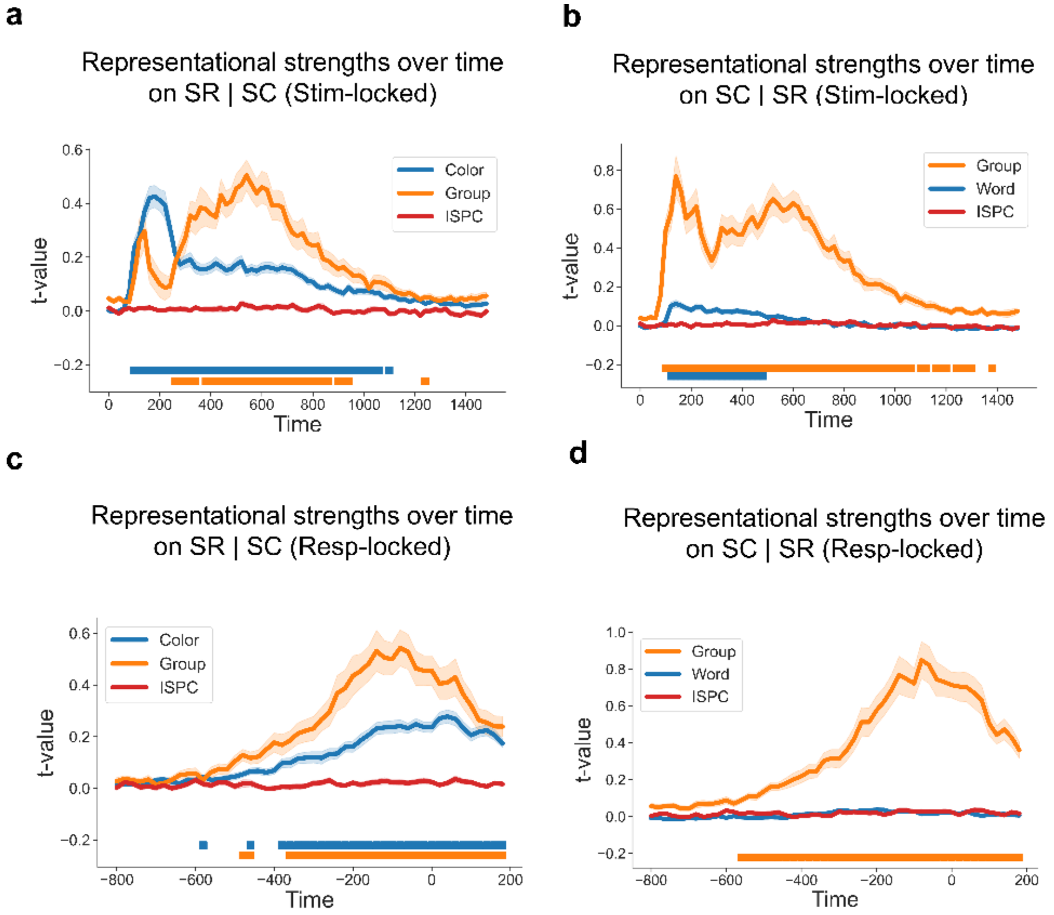


**Supplementary Figure 4 | Shared dimensions between SC and SR subspaces.** (**a**) Group average t values of representational strength for color, group and ISPC over time on SC decoder trained on SR subspace (SR | SC) with stimulus-locked analysis. Squares below the lines indicate the significant time points ($p < 0.001$, Bonferroni corrected). (**b**) Group average t values of representational strength for color, group and ISPC over time on SR decoder trained on SC subspace (SC | SR) with stimulus-locked analysis. Squares below the lines indicate the significant time points ($p < 0.001$, Bonferroni corrected). (**c**) Group average t values of representational strength for color, group and ISPC over time on SC decoder trained on SR subspace (SR | SC) with response-locked analysis. Squares below the lines indicate the significant time points ($p < 0.001$, Bonferroni corrected). (**d**) Group average t values of representational strength for color, group and ISPC over time on SR decoder trained on SC subspace (SC | SR) with response-locked analysis. Squares below the lines indicate the significant time points ($p < 0.001$, Bonferroni corrected).

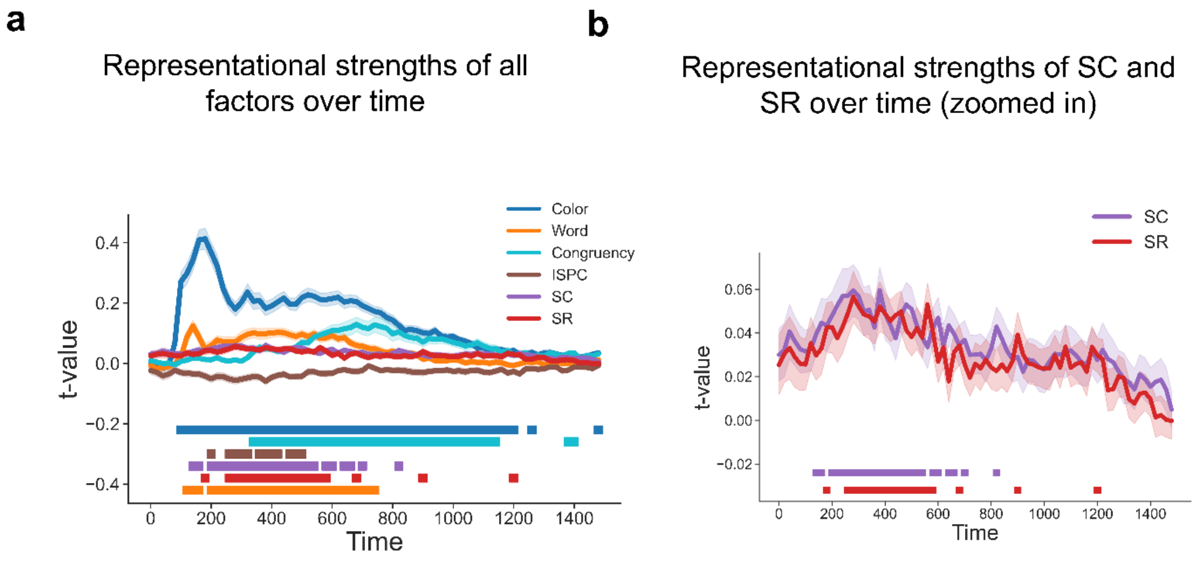


**Supplementary Figure 5 | Simultaneous EEG representations of SC and SR associations.** (**a**) Group average t values of representational strength for each factor over time. Squares below the lines indicate the significant time points ($p < 0.05$, Bonferroni corrected). (**b**) SC and SR association results from Supplementary Fig. 5a. Shaded areas denote SEM.

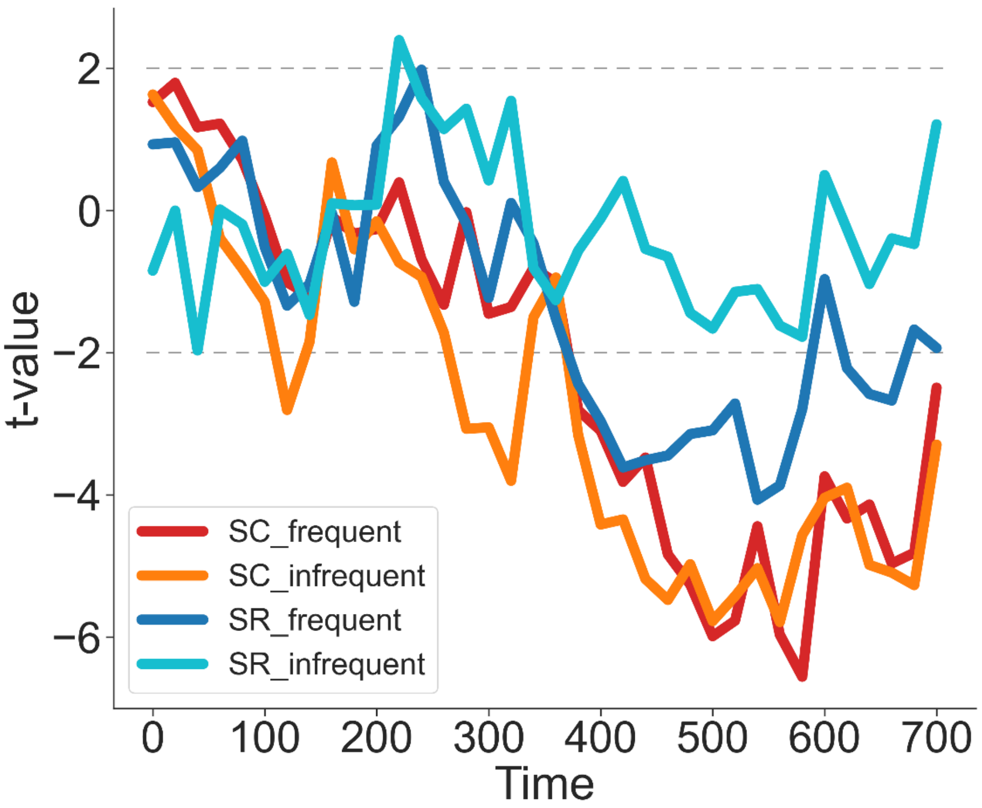


**Supplementary Figure 7 | The predictions of SC and SR effects on the RT on frequent and infrequent trials.** Group average *t* values of each factor predicting RT in the LMM analysis.

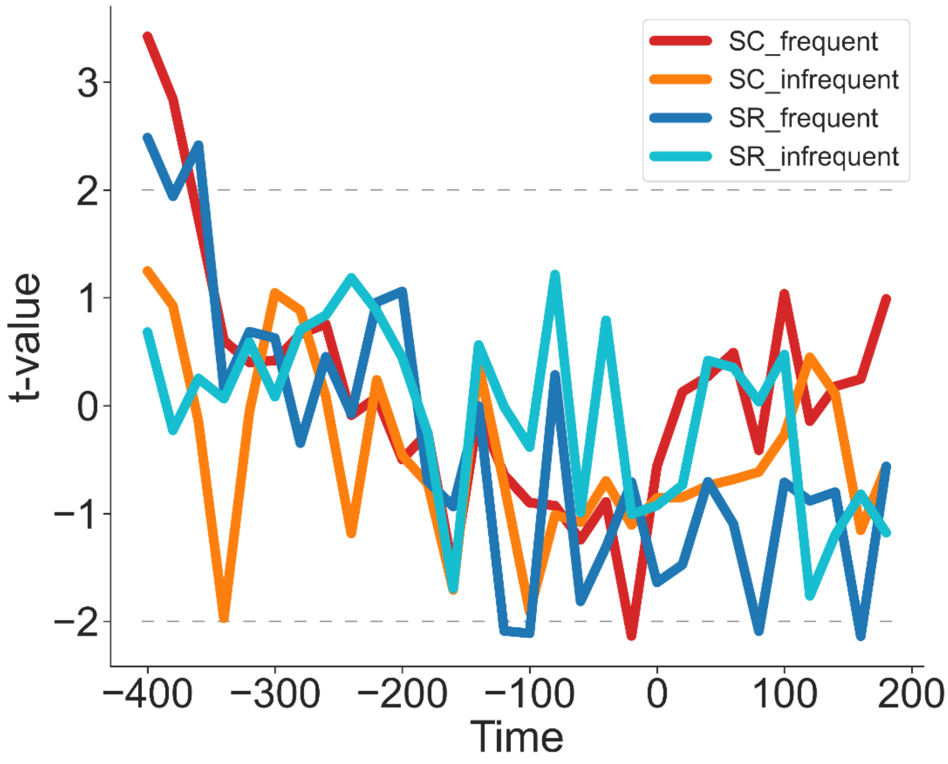


**Supplementary Figure 8 | The predictions of SC and SR effects on the RT on frequent and infrequent trials using response-locked analysis.** Group average *t* values of each factor predicting RT in the LMM analysis.

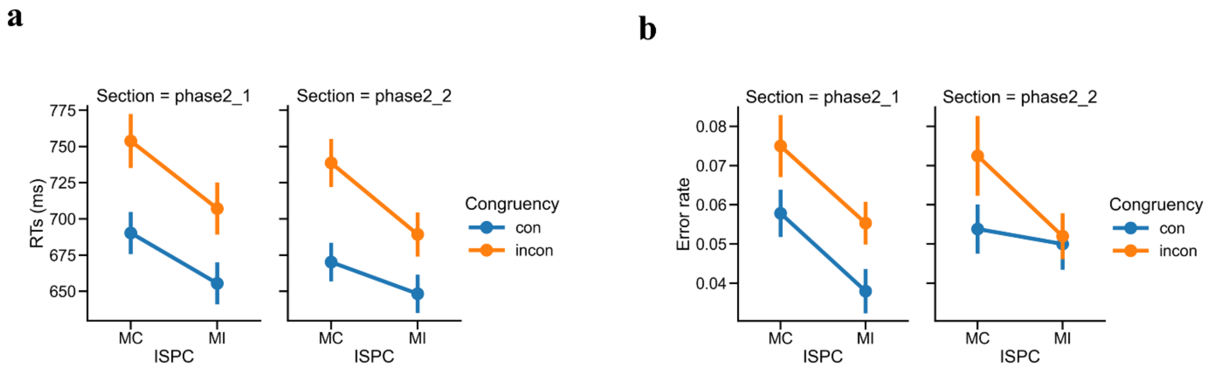


**Supplementary Figure 9 | ISPC effect in the first and second halves of Phase 2 (a)** Group mean error rate as a function of the ISPC effect. Error bars show standard errors of the mean (SEM). MI: mostly incongruent trials; MC: mostly congruent trials; incon: incongruent trials; con: congruent trials. **(a)** Group mean error rate as a function of the ISPC effect. Error bars show standard errors of the mean (SEM). MI: mostly incongruent trials; MC: mostly congruent trials; incon: incongruent trials; con: congruent trials.
